# Unique repression domains of Pumilio utilize deadenylation and decapping factors to accelerate destruction of target mRNAs

**DOI:** 10.1101/802835

**Authors:** René M. Arvola, Chung-Te Chang, Joseph P. Buytendorp, Yevgen Levdansky, Eugene Valkov, Peter L. Freddolino, Aaron C. Goldstrohm

## Abstract

Pumilio is an RNA-binding protein that represses a network of mRNAs to control embryogenesis, stem cell fate, fertility, and neurological functions in *Drosophila*. We sought to identify the mechanism of Pumilio-mediated repression and find that it accelerates degradation of target mRNAs, mediated by three N-terminal Repression Domains (RDs), which are unique to Pumilio orthologs. We show that the repressive activities of the Pumilio RDs depend on specific subunits of the Ccr4-Not (CNOT) deadenylase complex. Depletion of Pop2, Not1, Not2, or Not3 subunits alleviates Pumilio RD-mediated repression of protein expression and mRNA decay, whereas depletion of other CNOT components had little or no effect. Moreover, the catalytic activity of Pop2 deadenylase is important for Pumilio RD activity. Further, we show that the Pumilio RDs directly bind to the CNOT complex. We also report that the decapping enzyme, Dcp2, participates in repression by the N-terminus of Pumilio. These results support a model wherein Pumilio utilizes CNOT deadenylase and decapping complexes to accelerate destruction of target mRNAs. Because the N-terminal RDs are conserved in mammalian Pumilio orthologs, the results of this work broadly enhance our understanding of Pumilio function and roles in diseases including cancer, neurodegeneration, and epilepsy.

## INTRODUCTION

Proper control of gene expression is accomplished in part by RNA-binding proteins (RBPs) that affect processing, transport, translation, and degradation of messenger RNAs (mRNAs). *Drosophila* Pumilio (Pum) is a quintessential sequence-specific RBP that regulates the fate of mRNAs in the cytoplasm. Pum is a member of the eukaryotic PUF family (named after Pum and *C. elegans* fem 3-binding factor), which share a conserved Pum homology domain (Pum-HD) (1). Pum is essential for development and impacts a wide range of biological processes (2). Pum is broadly expressed and is abundant in embryos, the nervous system, and the female germline. During early embryogenesis, Pum represses expression of the morphogen Hunchback, a crucial factor in the establishment of polarity and body plan (3–9). In the germline, Pum regulates stem cell proliferation and differentiation (10–14). Moreover, Pum plays multiple roles in the nervous system, where it controls neuronal morphology, electrophysiology, motor function, and learning and memory formation (15–21).

Pum regulates specific mRNAs by binding to a short RNA sequence, 5’ UGUANAUA, termed the Pumilio Response Element (PRE), via its RNA-binding domain (RBD) that encompasses the Pum-HD and flanking residues (2,5,22–25). The RBD is comprised of eight repeats of a triple alpha-helical motif which form an arched molecule that recognizes single-stranded RNA (25, 26). Each repeat presents three amino acids that specifically interact with a ribonucleotide base. Pum binds to an extensive network of mRNAs, the majority of which contain one or more PREs located in the 3’ untranslated region (3’UTR) (2,5,27–29).

Notwithstanding substantial insights into Pum’s biological roles, structure, and RNA-binding activity (2), our understanding of the mechanisms by which it represses gene expression remains incomplete. An early model proposed that Pum recruits Nanos (Nos) and Brain tumor (Brat) to block translation of *hunchback* mRNA (30–32); however, recent developments have substantially revised that model. We now know that Pum, Nos, and Brat are each sequence specific RBPs that can combinatorially regulate a subset of mRNAs (2,25,28,33,34). Nos can bind in a cooperative manner with Pum to certain mRNAs that contain a Nos Binding Site (NBS) immediately upstream of a PRE, thereby strengthening Pum-mediated repression (25). Additionally, Brat was shown to bind specific mRNAs on its own and confers repressive activity independent of Nos or Pum (28,33,34). In the case of the *hunchback* mRNA in embryos, Brat, Pum, and Nos collectively repress it by binding to two Nos Response Elements (NREs), each of which contain a Brat binding site, an NBS, and a PRE (2,25,28,33–35). Importantly, Pum can repress PRE-containing mRNAs independent of Nos or Brat (36). For example, Pum potently represses PRE-bearing reporter mRNAs in cultured *Drosophila* d.mel2 cells that do not express detectable Nos. Moreover, depletion of Nos and/or Brat did not alter Pum’s ability to repress. Further, Pum can repress mRNAs that are not bound by Nos or Brat. In this study, we focus on determining the mechanism by which Pum represses mRNAs. The resulting knowledge will be essential to understand how Pum regulates its multitude of targets and how it collaborates with other RBPs, such a Nos and Brat, to regulate subsets of those mRNAs.

Multiple studies have provided insights into the mechanism of Pum-mediated repression. Early evidence correlated repression of *hunchback* mRNA by Pum - along with Nos and Brat – during embryogenesis with shortening of that transcript’s 3’ poly-adenosine (poly(A)) tail (i.e. deadenylation) (8, 35). The poly(A) tail promotes translation and stability of mRNAs, and deadenylation reduces protein expression and initiates mRNA decay (37, 38). Like all eukaryotes, *Drosophila* possesses multiple deadenylase enzymes (39–41). Pum was reported to interact with the Ccr4-Not (CNOT) complex (42–44), which contains both Pop2/Caf1 and Ccr4/twin deadenylases.

Pum also cooperates with Nos or Brat in other contexts, and again deadenylation is implicated. In the germline, Pum and Nos regulate *cyclin B* (*cycB*) in pole cells and *mei-P26* mRNA in germline stem cells (GSCs) (42, 43). In both cases, Pum and Nos are thought to utilize the CNOT deadenylase complex. Pum and Brat regulate targets in the cystoblast to attenuate the local effects of Dpp signaling, and this effect is thought to require CNOT, as the Pop2 deadenylase was necessary for Pum and Brat to repress a reporter bearing the *mad* 3’UTR (11). In terms of the Pum repression mechanism, a complication in interpreting these experiments is that Nos and Brat are also linked to CNOT and deadenylation (40,45,46). Thus, it was necessary to develop approaches that specifically dissect repression of mRNAs by Pum alone.

We previously used PRE-containing reporter genes to measure Pum repression activity in *Drosophila* cells and showed that it reduces both protein and mRNA levels (36). Four regions of Pum contribute to its repressive activity. The highly conserved RBD made a minor contribution, whereas the N-terminus of Pum contains the major repressive activity. Repression by the Pum RBD required a poly(A) tract in the target mRNA and the cytoplasmic poly(A) binding protein (pAbp) (44). The Pum RBD associates with pAbp and antagonizes its ability to promote translation. The Pum RBD also interacts with Pop2 and promotes deadenylation (42, 44). While depletion of Pop2 and Ccr4 blocked mRNA decay induced by the Pum RBD, it did not prevent RBD-mediated translational repression, whereas pAbp depletion did. Thus, the Pum RBD appears to primarily act via inhibition of poly(A)-dependent translation.

The robust repressive activity of the Pum N-terminus is conferred by three repression domains (Figure 1A, RD1, RD2, and RD3) (36). These RDs are unique to Pum orthologs spanning from insects to vertebrates (47). They do not share homology with each other or previously characterized protein domains. Each is capable of repressing protein expression when directed to a reporter mRNA (36). The crucial remaining challenge is to determine how the Pum N-terminal RDs regulate target mRNAs. In this study, we characterize their regulatory activities and investigate the co-repressors necessary for repression.

**Figure 1.**
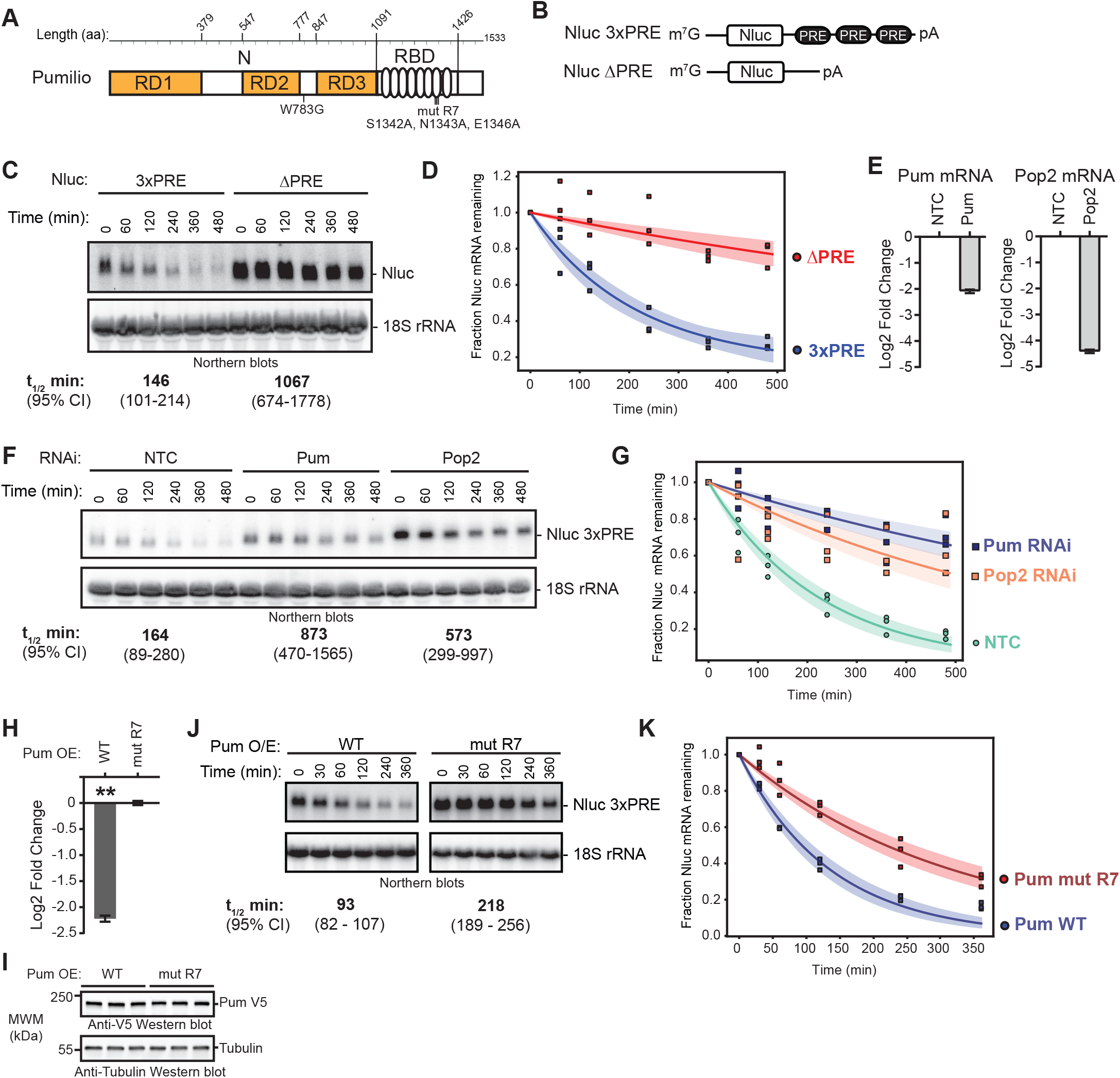
Pumilio accelerates mRNA degradation. (A) Diagram of *Drosophila melanogaster* Pumilio (Pum) protein with N-terminal (N) Repression Domains (RD1, RD2 and RD3) and C-terminal RNA-Binding Domain (RBD). Amino acid boundaries are listed at the top. Amino acid substitutions used in this study, including the putative cap-binding amino acid (W783G) and the RNA-binding defective mutant repeat seven (mut R7), are annotated below the diagram. (B) Diagram of Nano-luciferase (Nluc) reporter mRNAs containing three Pum Response Elements (3x PRE) sequences in 3’UTR, along with 7-methyl guanosine cap (m7G) and poly(A) tail (pA). An equivalent Nluc reporter lacking the PRE sequences (ΔPRE) was used as a control. Diagram is not to scale. (C) Transcription shut-off with Actinomycin D (ActD) was performed to compare the half-lives of the Nluc 3xPRE and Nluc ΔPRE reporter mRNAs in d.mel2 cells. A representative Northern blot of Nluc reporters and the 18S ribosomal rRNA internal control is shown. Each lane of the gel contains 10 µg of total RNA. The measured half-life of each reporter mRNA is shown below the respective blots. Mean values from three experimental replicates are reported with 95% credible intervals. Data and statistics are reported in Supplementary Table S1. (D) The fraction of Nluc mRNA remaining, normalized to internal control 18S rRNA, is plotted relative to time (minutes) after inhibition of transcription. Data points for each of three experimental replicate are plotted. First order exponential decay trend lines, calculated by non-linear regression analysis, are plotted for each experimental condition (dark lines), along with 95% credible interval for each trend line (shaded areas: orange for Nluc ΔPRE and blue for 3xPRE). (E) RNAi-mediated depletion of Pum or Pop2 mRNAs after 4 days of dsRNA treatment was measured by RT-qPCR. The Pum or Pop2 mRNA level was normalized to internal control Rpl32 mRNA and fold change was calculated relative to the non-targeting control RNAi condition (NTC). The mean log_2_ fold change of the indicated mRNA level is plotted with 95% credible intervals based on 3 biological replicates with three technical replicates each. Data and statistics are reported in Supplementary Table S1. (F) Transcription shut-off with ActD was performed using the Nluc 3xPRE reporter mRNA to measure the effect of RNAi depletion of Pum or Pop2 relative to NTC. A representative Northern blot of Nluc 3xPRE reporter and 18S rRNA is shown. Each lane of the gel contains 10 µg of total RNA. Half-lives of the mRNA in the respective conditions, determined from 3 experimental replicates, are shown below the diagram, along with 95% credible intervals. Data and statistics are reported in Supplementary Table S1. (G) The fraction of Nluc mRNA remaining, normalized to internal control 18S rRNA, is plotted relative to time (minutes), after inhibition of transcription. Data points for each of 3 experimental replicates are plotted. First order exponential decay trend lines, calculated by non-linear regression analysis, are plotted for each experimental condition (dark lines), along with 95% Credible Interval for each trend line (shaded areas: orange for RNAi of Pop2, blue for RNAi of Pum, and green for negative control RNAi, NTC). (H) Repression of Nluc 3xPRE reporter activity by wild type (WT) over-expressed (OE) Pum in d.mel2 cells was measured by dual luciferase assay. Nluc activity was normalized to Firefly luciferase expression from a co-transfected plasmid in each sample. Mean log_2_ fold change in normalized Nluc 3xPRE activity by WT Pum is plotted relative to the RNA-binding defective mutant Pum (mut R7) along with 95% credible intervals, as determined from four technical replicate measurements from three biological replicate samples. Data and statistics are reported in Supplementary Table S1. For significance calling, the ‘**’ indicates a posterior probability of >0.95 that the indicated difference is at least 1.3-fold. (I) Western blot detection of over-expressed, V5 epitope-tagged WT or mut R7 Pum in three biological replicate samples each. Each lane contains an equivalent mass of cell extract, as measured by Lowry assay. Western blot of Tubulin served as a control for equivalent loading of the samples. (J) Transcription shut-off with ActD was performed to measure half-life of Nluc 3x PRE reporter in response to over-expressed wild type Pum or mut R7. Northern blots of Nluc 3x PRE reporter and 18S rRNA from a representative experiment are shown and half-lives and 95% credible intervals from 3 biological replicates are reported at the bottom. Each lane of the gel contains 10 µg of total RNA. Data and statistics are reported in Supplementary Table S1. (K) The fraction of Nluc mRNA remaining, normalized to internal control 18S rRNA, is plotted relative to time (minutes) after inhibition of transcription. Data points for each of 3 biological replicates are plotted. First order exponential decay trend lines, calculated by non-linear regression analysis, are plotted for each experimental condition (dark lines), along with 95% credible interval for each trend line (shaded areas: orange for Pum mut R7, blue for Pum WT). We note that the last time point (360 minutes) for both conditions was excluded from the analysis because signs of saturation/plateau were apparent for the WT case; thus, data from that time point are plotted, but not included in the fit.

An earlier model proposed that the 5’ 7-methylguanosine cap of target mRNAs is important for Pum repression (48). The 5’ cap plays a key role in translation and mRNA stability, and its enzymatic removal (i.e. decapping) initiates 5’ mRNA decay (37, 38). Analysis of the *Xenopus* Pum ortholog identified a 5’ cap-binding motif that contributes to cap-dependent translation inhibition in oocytes (48). Because this motif is conserved in *Drosophila* Pum (11, 48), it was postulated to contribute to translational inhibition; however, conflicting data have been reported. First, deletion of the putative cap-binding region (PCMb) did not alleviate Pum repression, and the cap-binding region did not display repression activity when directly tethered to an mRNA (36). In contrast, another study reported that mutation of a conserved tryptophan in the cap-binding motif reduced Pum activity (11). In this study, we further scrutinize the potential contribution of this motif.

A previous study reported repression by Pum and Nos of reporter genes whose translation is driven by either 5’ cap-dependent or independent translation using a *Drosophila* eye phenotypic assay (49). There are several important caveats to that analysis. First, the internal ribosome entry site that was used remains poorly characterized (50, 51). Second, the contribution of mRNA decay was not assessed. Third, the experimental system could not separately analyze contributions by Pum and Nos. Therefore, the potential relevance remains unknown.

In this report, we show that Pum accelerates mRNA degradation of PRE-bearing mRNAs, mediated by N-terminal repression domains. We find that the putative 5’ cap binding motif is not necessary for Pum repression. Instead, the Pum RDs directly bind to the CNOT deadenylase complex, and specific CNOT subunits are required to repress and degrade target mRNAs. We also detect that the Pum N-terminus has an additional repressive activity that circumvents the requirement for CNOT and the poly(A) tail and involves the mRNA decapping enzyme, Dcp2. We measured the contribution of multiple mechanisms to repression by Pum, emphasizing the importance of CNOT, Dcp2, and pAbp. Taken together, our data reveals that Pum utilizes deadenylation and decapping pathways to repress and degrade its target mRNAs.

## MATERIALS AND METHODS

### Plasmids and cloning

The plasmids used in this study are listed in **Supplementary File 1** and were verified by DNA sequencing. The sequences of all oligonucleotide primers are listed in **Supplementary File 1**. The pIZ plasmid (Invitrogen) was used for effector expression and contains the OpIE2 promoter, *Drosophila* Kozak sequence, C-terminal V5 epitope and His6 tags, and the SV40 cleavage/poly-adenylation site. For experiments employing over-expression of Pumilio, the coding region of wild type (NP_001262403.1) or RNA-binding defective (mut R7) Pumilio was cloned into pIZ to create pIZ Pumilio V5H6 and pIZ Pumilio mut R7 V5H6 plasmids, as previously described (36). Amino acid residues S1342A, N1343A, and E1346A of the seventh repeat of the Pum-HD are mutated in the RNA-binding defective Pum mut R7 (Figure 1A), as previously described (25, 36).

For tethered function assays, the MS2 fusion effector-encoding plasmids pIZ MS2-PumN V5H6, pIZ MS2-RD1 V5H6, pIZ MS2-RD2 V5H6, and pIZ MS2-RD3 V5H6 plasmids were previously described (36). The negative control pIZ MS2-EGFP plasmid was created by inserting the coding sequence for enhanced green fluorescent protein (EGFP), amplified by PCR using oligos CW115 and CW116, into KpnI and XbaI sites of pIZ MS2CP vector (36). Likewise, the Dcp1 coding sequence (NP_611842.1, amplified with oligos RA 102 and RA 103) was inserted into SpeI and NotI sites to create pIZ MS2-Dcp1.

For RNAi rescue experiments, cDNA clone pIZ myc-Pop2 was generated by insertion of the Pop2 coding sequence (NP_648538.1, amplified using oligo CW 033 and CW 034) with an N-terminal Myc tag into HindIII and XbaI sites of the pIZ plasmid. Pop2 mutations, D52A and E54A, were introduced into pIZ myc-Pop2 using quickchange site-directed mutagenesis (Agilent) with primers CW 161 and CW 162.

The expression plasmid vector pUBKz 3x Flag contains the *Drosophila* ubiquitin 63E promoter, Kozak sequence, N-terminal 3x Flag, and SV40 cleavage and poly-adenylation site in a pUC19 backbone (provided by Dr. Eric Wagner, University of Texas Medical Branch). For inhibition of decapping by over-expression, the plasmid pUbKz 3x Flag Dcp2 E361Q was created by inserting the Dcp2 coding sequence (NP_001246776.1, amplified with oligos RA 086 and RA 087) into SpeI and NotI sites in pUbKz 3x Flag vector followed by site directed mutagenesis to introduce the E361Q mutation (as used in (45,52,53)) using oligos RA 166 and RA 167.

Reporter genes are based on vector pAc5 (Invitrogen). The internal control plasmid, pAc5.1 FFluc min 3’UTR, which expresses firefly luciferase, was described previously (36). The reporter plasmid pAc5.4 Nluc2 ΔPRE 3’UTR was cloned by inserting the Nano-luciferase (Nluc) coding sequence with C-terminal PEST sequence, derived from pNL1.2 plasmid (Promega) and amplified using oligos CW 578 and RA 066, into the KpnI and XhoI sites of vector pAc5.4 (25). The tethered function reporter plasmid pAc5.4 Nluc2 2xMS2 was cloned by inserting oligos AG 784 and AG 785, encoding two copies of the binding site for MS2 coat protein (MS2), into Xho1 and Not1 sites of pAc5.4 Nluc MCS. To create the Histone Stem Loop (HSL) reporter, pAc5.4 Nluc2 2xMS2 HSL, inverse PCR with oligos RA 255 and RA 256 was performed using pAc5.4 Nluc2 2xMS2 template, thereby replacing cleavage/poly-adenylation element with the HSL and Histone Downstream Element (HDE) sequences. To create the Pum reporter plasmid, first a unique XhoI site was inserted into pAc5.1 Rnluc (36) using inverse PCR with oligos RA 214 and RA 215. Next, the Nluc2 coding sequence was inserted into KpnI and XhoI sites in the pAc5.1 vector to create pAc5.1 Nluc2 3xPRE.

For production of recombinant Pum constructs in *E. coli*, cDNA sequences encoding Pum RD1 (aa 1-378), RD2 (aa 548-776), RD3 (aa 848-1090), or Pum RBD (aa 1091-1426) were inserted using the Gibson assembly method (54) into the pnYC-pM plasmid vector (55) linearized with NdeI. The resulting Pum RD1, RD2, RD3, and RBP fusion proteins have N-terminal MBP tags that are cleavable by the human rhinovirus 3C (HRV3C) protease, and a C-terminal StrepII tag.

### Cell Culture and Transfection

D.mel-2 cells (Invitrogen) were cultured in Sf900III media (Thermo-Fisher, see **Supplementary File 1** for reagents) at 25°C in 100 µg/ml penicillin and 100 µg/ml streptomycin. In our standard transfection procedure, 2 million d.mel-2 cells were plated in a 6-well plate and transfected with 150 µl of transfection mix containing FuGene HD (Promega) and 3 µg effector DNA at a 4 µl Fugene HD:1 µg DNA ratio in Sf900III media. This transfection mix was incubated for up to 15 minutes at room temperature prior to application to cells. In experiments that measure regulation by endogenous Pum, the transfection mix contained a total of 1.5 µg of transfected DNA with FuGene HD. For dual luciferase assays and Northern blotting, 20 ng of pAc5.4 NLuc poly(A) or 100 ng Nluc HSL, along with 20 ng pAc5.1 Ffluc, were included in the transfection mix.

For the Pop2 rescue experiments, 5 ng of pAc5.4 Nluc 2xMS2 poly(A) reporter and pAc5.1 FFluc internal control plasmid were included in the transfection mix. In the Pop2 RNAi rescue experiment (Figure 4F-4H), cells were transfected with 2.25 µg of the indicated tethered effector and 750 ng of either pIZ EGFP V5, pIZ myc-Pop2, or pIZ myc-Pop2 D52A E54A. For analysis of Not1 RNAi rescue with exogenous Pop2 (Supplementary Figure S3), either 150 ng of pIZ EGFP V5 control, 100 ng myc-Pop2 (with 50 ng pIZ EGFP V5), or 150 ng myc-Pop2 were transfected into cells, along with 2.85 µg of the indicated MS2-tethered effectors.

For analysis of decapping, cells were transfected with 1.5 µg of the indicated MS2-tethered effectors and 1.5 µg of either pUbKz 3x Flag empty vector or pUbKz 3x Flag Dcp2 E361Q plasmid. In this approach, the cells were also treated with either non-targeting control (NTC) or Dcp2 double stranded RNA, as indicated in the figure. Importantly, RNAi of the Dcp2 mRNA targeted the 3′UTR and thus did not affect expression of Dcp2 E361Q. To measure regulation by endogenous Pum and co-repressors in Figure 11, the transfection mix contained a total of µg of transfected DNA with FuGene HD, along with 20 ng of FFluc and 20 ng of the indicated Nluc reporter plasmids.

### RNA interference

To induce RNAi, gene-specific double stranded RNA (dsRNA), ranging from 133-601 bp, were designed using the SnapDragon web-based tool provided by the Harvard Drosophila RNAi Screening Center (URL: http://www.flyrnai.org/cgi-bin/RNAi_find_primers.pl) to minimize potential off-target regions. Transcription templates were PCR-amplified with primers that add opposing T7 promoters to each DNA strand (see **Supplementary File 1** for dsRNA template primer sequences with T7 RNA polymerase promoters). The dsRNAs were transcribed from these templates using HiScribe T7 high yield RNA synthesis kit (New England Biolabs). The dsRNAs were then treated with RQ1 RNase-free DNase (Promega) and purified using RNA Clean & Concentrator-25 (Zymo Research). Non-targeting control (NTC) dsRNA, corresponding to the *E. coli* LacZ gene, was described previously (36).

For all RNAi experiments measuring dual luciferase activity, d.mel-2 cells were plated in 6-well plates with 24 µg dsRNA per well. In the standard protocol, one million cells were plated with dsRNA, incubated 24 hours, then reporters and effectors were transfected with FuGene HD as described above. Forty-eight hours after transfection cells were harvested for luciferase assays, Western blotting, and RNA isolation. For Figure 4F-H, Figure 10 and Supplementary Figure 3, a half-million cells were plated and incubated with dsRNAs for 72 hours, then reporters and effectors were transfected with FuGene HD as described above. Forty-eight hours after transfection, cells were harvested for luciferase assay, Western blotting and RNA isolation.

### Reporter gene assays

D.mel-2 cells were harvested from a transfected 6-well plate and 100 µl of cell culture (∼0.5-6 x10^5^ cells, depending on experimental conditions) was aliquoted into a 96-well plate. Luciferase assays were then performed using Nano-Glo Dual-Luciferase Reporter Assay System (Promega) and a GloMax Discover luminometer (Promega) per manufacturer’s instructions, using 10 µl/ml of Nluc substrate. To measure regulation by endogenous or over-expressed Pum, the Nluc 3xPRE reporter was used. To measure activity of tethered effectors, the Nluc 2xMS2 poly(A) or Nluc 2xMS2 HSL reporters were used. The internal control pAC5.1 FFluc 3’UTR poly(A) was used in all reporter experiments. In tethered assays, MS2-EGFP served as the negative control for effectors. In experiments analyzing repression by full-length wild type or mutant W783G Pum, the RNA-binding defective mutant Pum (mut R7) served as a negative control, as previously established (36).

### Data and Statistical Analyses

For reporter gene assays, the Nluc and FFluc reporter activities from each sample, measured in Relative Light Units (RLU), were used to calculate fold change response values. First, Relative Response Ratios (RRR) for each sample were calculated by dividing the Nluc value by the FFluc value to normalize variation in transfection efficiency. Next, the fold change in RRR for a given effector/condition was calculated relative to a negative control effector/condition. For tethered function assays, fold change by an effector was determined relative to the mean RRR for the negative control effector, MS2-EGFP, unless otherwise noted. For tethered function assays utilizing RNAi of a putative co-repressor, fold change induced by an effector was measured relative to the negative control effector, MS2-EGFP, within the same RNAi condition.

For Pum-mediated repression of the PRE-containing reporter, Nluc 3xPRE, the fold change induced by wild type or mutant Pum was determined relative to the mean RRR for the RNA-binding defective Pum mut R7 negative control. For RNAi experiments in Figure 11C, fold change induced by depletion of a regulatory factor was determined relative to the mean RRR for the non-targeting control siRNA. The data was analyzed by determining the RRR of Pum-repressed Nluc 3xPRE reporter activity, normalized to the RRR of the non-Pum regulated Nluc ΔPRE reporter activity within the same RNAi condition. In doing so, the Pum specific effect is measured while normalizing for Pum/PRE independent effects of the RNAi. From this data, the fold change in reporter expression induced by each RNAi condition was calculated relative to the NTC. The data were separately analyzed and reported in Figure 11D in the same manner, except that the FFluc values were omitted from the calculations. The PRE-dependent effect of each RNAi condition on the Nluc 3xPRE reporter was normalized to the effect on the unregulated Nluc ΔPRE reporter. The fold change in PRE-mediated regulation within each RNAi condition was then calculated relative to the negative control NTC dsRNA.

To assess replication and reproducibility of measurements, we employed several types of replicates including experimental replicates (i.e. independent assays performed on separate days using the same approach), biological replicates (i.e. parallel measurements of distinct cell samples) and technical replicates (i.e. multiple measurements from the same sample). The number and type of replicates for each experiment are indicated in the figure legends and data tables (**Supplementary Table 1**), and were dictated by the assay type, experimental design, and feasibility.

For statistical analysis of reporter gene data, a hierarchical Bayesian model was used to account for the differences in variation between technical, biological, and experimental replicates. The input data were the ratios of Nluc activity to that of the FFluc internal control. The average value for each replicate was modeled as arising from a t-distribution centered on the overall average value µprot for the particular effector protein; a separately inferred average value for the Ffluc intensity on each particular day (µFfluc,day, shared across all effector proteins considered on that day) was applied as an additive offset to the day-wise values. The values for the technical replicates were further modeled with t-distributions centered on the appropriate daily average value µprot,day - µFfluc,day. The protein-wise scale parameters µprot were modeled as arising from a common gamma distribution with uninformative hyperpriors, whereas the other scale parameters, and the degree of freedom parameters for the t distributions were simply assigned uninformative priors. We fitted the models using JAGS via the rjags interface, running four independent Monte Carlo chains for 250,000 iterations with 25,000 steps of burn-in; we ensured convergence by verifying that the Gelman-Rubin shrinkage statistic for all parameters of interest was <1.1. Reported credible intervals were calculated using the highest posterior density approach with the R boa package; reported probabilities were calculated directly from the posterior distributions. We performed separate fits for each reporter construct. The main topic of interest in all cases is the central log ratio value µprot for each protein, which reflects the relative Luciferase level observed when that protein is present in the assay. We most frequently report the differences observed for the µprot value of a protein effector relative to that of the negative control.

All data, number and types of replicates, and statistics (e.g. credible intervals and posterior probabilities) from the fitted models are reported in **Supplementary Table 1**. Unless otherwise noted, graphs represent mean fold change values and 95% credible intervals, as described in the figure legends.

### Pum antibody

The anti-Pumilio rabbit polyclonal antibody was generated using the recombinant purified antigen containing Pum residues 1434-1533 fused to GST. Pum-specific antibodies were antigen-affinity purified from the resulting serum using a column containing immobilized, recombinant, purified Halotag-Pum aa1434-1533 immobilized to Halolink resin (Promega) (56). The Pum antigen sequence was: PITVGTGAGGVPAASSAAAVSSGATSASVTACTSGSSTTTTSTTNSLASPTICSVQENGSAMVVEPSSPDASESSSSVVSGAVNSSLGPIGPPTNGNVVL.

### Western blotting

Cell lysates were prepared by adding 100 µl of lysis buffer (50 mM Tris-HCl pH 8.0, 150 mM NaCl, 2x complete mini, ethylenediaminetetraacetic acid (EDTA)-free protease inhibitor cocktail (Roche), and 0.5% Non-Idet P40 (NP40)) to cell pellets containing 0.5-6 x10^6^ cells, dependent on experimental design. Cells were lysed for 10 seconds using a cell disruptor. Cell debris was removed by centrifugation at 21,100 x g for 10 minutes. Total protein in the resulting cell lysate was quantitated using Lowry DC assay (BioRad) with a bovine serum albumin (BSA) standard curve. Equal mass (10 µg, unless noted otherwise) of cell lysates were then analyzed on SDS-PAGE gels (4–20% Mini-PROTEAN TGX or Criterion TGX, BioRad) along with PageRuler Prestained Plus Molecular Weight Markers (Thermo Fisher). For detection of endogenous Pum protein, 30 µg of total cellular protein per lane was analyzed. Protein was then transferred onto Millipore Immobilon-P (or Immobilon-PSQ for detection of Pop2 and EGFP) polyvinylidene difluoride (PVDF) membrane at 30 V overnight or 65 V for 1.5 hours. Pop2 and Ccr4 western blots were blocked in tris-buffered saline (TBS: 20 mM Tris-HCl pH 7.5, 137 mM NaCl, 0.1% Tween 20) with 3% BSA, and TBS+BSA was used in all subsequent steps. All other blots were blocked with Blotto (5% powered dry nonfat milk in 1x phosphate-buffered Saline (10 mM Na_2_HPO_4_, 1.8 mM KH_2_PO_4_ pH 7.4, 2.7 mM KCl and 137 mM NaCl and 0.1% Tween 20).

Primary antibodies used in this study, and the working dilutions, are indicated in **Supplementary File 1**, and were incubated for either 1 hour at room temperature or overnight at 4°C. Blots were then washed three times with Blotto or TBS+BSA for 5 minutes per wash. The appropriate secondary antibody-horse radish peroxidase conjugate was the added at dilutions indicated in **Supplementary File 1,** and incubated for 1 hour at room temperature. The blots were then washed an additional three times. Blots were then incubated with either Pierce or Immobilon enhanced chemiluminescent (ECL) substrates for 1 min followed by colorimetric and chemiluminescent detection using a ChemiDoc Touch imaging system (BioRad). Western blot images were processed using Image Lab 5.2.1 software (BioRad). Images were exported to TIF files and processed for figures using Adobe Creative Suite. In figures, the western blot images from the same antibody, blot and exposure are surrounded by black boxes. In the event that lanes were cropped from the same blot image, white space is made apparent.

### Immunoprecipitation

D.mel-2 cells (2 million per sample) were transfected with the Flag-tagged bait protein and V5-tagged prey protein expression plasmids indicated in Figure 7 using FuGene HD as described above and incubated for 3 days at to allow protein expression. Cells were lysed with a cell disruptor in Flag Buffer A (50 mM Tris-HCl pH 8.1, 200 mM NaCl, 1 mM EDTA, 0.2% Triton X100 and 2x complete protease inhibitor cocktail (Roche) and cell debris was removed by centrifugation at 21,100 x g for 10 minutes. Cell extracts were split into RNase treated (4 units RNase One, Promega) and untreated (120 units of RNasin ribonuclease inhibitor, Promega) samples and incubated with 10 µl bed volume of EZview Red anti-FLAG M2 Affinity Gel (Sigma) overnight at 4°C. Beads were washed three times with Flag Buffer A, and three times with Flag Buffer B (50 mM Tris-HCl pH 8.1, 200 mM NaCl, and 1 mM EDTA) and then resuspended in 60 µl Flag Buffer B. Bound proteins were eluted by heating in 1 x SDS-PAGE loading dye. Samples were then analyzed by SDS-PAGE and western blot.

### In vitro pulldown assays

The StrepII-tagged MBP and MBP-tagged Pum fragments (RD1, RD2, RD3, and RBD) were expressed in E. coli BL21 (DE3) Star cells (Thermo Fisher Scientific) grown in LB medium overnight at 37°C. Cells were lysed in lysis buffer (8 mM Na_2_HPO_4_, 137 mM NaCl, 2 mM KH_2_PO_4_, 2.7 mM KCl, 0.3% (v/v) Tween-20, pH 7.4). The cleared lysates were incubated with 30 µl (50% slurry) of StrepTactin sepharose resin (IBA). After 1 hour incubation, the beads were washed three times with lysis buffer and once with binding buffer (50 mM Tris-HCl pH 7.5, 150 mM NaCl). Purified human CCR4–NOT complex (50 µg) was then added to the beads. The reconstitution of the human CCR4–NOT complex was described elsewhere (57) and includes the following eight components: CNOT1 (amino acids 1-2376), CNOT2 (1–540), CNOT3 (1–753), CNOT10 (25–707), CNOT11 (257–498), CAF1 (1–285), CCR4a (1–558), CAF40 (1–299). After 1 hour incubation, the beads were washed three times with binding buffer and the proteins were eluted with binding buffer supplemented with 2.5 mM D-desthiobiotin. The eluted proteins were analyzed by SDS-PAGE and Coomassie blue staining. Pulldown results were confirmed in three independent experiments.

### Transcription shut off

To measure mRNA decay rates, transcription was shut off using Actinomycin D (58). To measure mRNA decay rates under endogenous Pum expression in Figure 1, 7.9 million d.mel-2 cells were seeded in a T75 flask containing 15.8 ml of Sf900III media. The cells were treated with a final concentration of 12 µg/ml of the indicated dsRNA and transfected with reporter plasmid 1 day after being seeded. Reporter plasmids were transfected into cells using FuGene HD as described above (scaled proportionally from 6-well format by surface area) with 23.7 µg of pIZ EGFP and 158 ng of pAc5.4 Nluc 3xNRE or ΔPRE reporter.

For analysis of over-expressed wild type or mutant mut R7 Pumilio on mRNA decay (Figures 1 and 6), 15.8 million d.mel-2 cells were seeded in a T75 flask containing 15.8 ml of Sf900III media and then were transfected with reporter plasmid. In the RNAi experiments in Figure 6, the cells were treated with a final concentration of 12 µg/ml of the indicated dsRNA immediately before transfection. Reporter plasmids were transfected into cells using FuGene HD as described above with 23.7 µg of pIZ Pum WT or mut R7 and 158 ng of pAc5.4 Nluc 3xNRE reporter. Three days post-transfection, transcription was inhibited by addition of Actinomycin D (Sigma) at a final concentration of 5 µg/ml. Prior to drug addition, two milliliters of cell culture was harvested (T=0 minutes). RNA was then purified from cells collected at time points including 2.5 ml of cell culture at each indicated time point, and 3.6 ml of cell culture at the final time point indicated in the corresponding figure.

### RNA purification and Northern blotting

RNA was isolated from d.mel-2 cells using the SimplyRNA Cells Low Elution Volume kit and Maxwell 16 RSC instrument (Promega). The RNA was quantitated using a NanoDrop spectrophotometer (Thermo Scientific) and its integrity was assessed by gel electrophoresis. For Northern blotting, total RNA (5 or 10 µg, as indicated in figure legends) was combined with 0.04 µg/µl Ethidium Bromide in sample buffer (23% formamide, 3% formaldehyde, 4.6 mM MOPS (3-(N-morpholino)propanesulfonic acid) pH 7, 1.1 mM sodium acetate, and 0.2 mM EDTA), and loading dye (2.1% glycerol, 4.2 mM EDTA, and 0.01% Bromophenol Blue and Xylene Cyanol) and heated at 75°C for 10 minutes. RNA was electrophoresed through a 1% denaturing agarose gel containing 1.48% formaldehyde and 1x MOPS buffer (20 mM MOPS pH 7.0, 5 mM sodium acetate, and 1 mM EDTA). The gels were imaged using UV detection with a ChemiDoc (BioRad) prior to transfer to assess integrity, migration of the ribosomal RNA (rRNA) and equivalent loading of lanes. The RNA was then blotted onto Immobilon-Ny+ membrane (Millipore) overnight using capillary transfer in 20x SSC buffer (3 M NaCl and 300 mM Sodium citrate), as previously described (25). The blot was then crosslinked with 120 J/cm^2^ UV (λ=254 nm) using a CL-1000 crosslinker (UVP). The blot was then either probed immediately or stored at 4°C.

Radioactive antisense RNA or DNA probes were used for Northern blot detection. Transcription templates for Nluc and FFluc antisense RNA probes were PCR-amplified using DNA oligonucleotides with a T7 RNA polymerase promoter appended to the antisense strand, described in **Supplementary File 1**. Using these templates, in vitro transcription was performed for 10 min at 37°C with the T7 MAXIscript transcription kit (Thermo-Fisher) in the supplied 1x Transcription Buffer with 1 µg of DNA template, 0.4 mM final concentration of ATP, CTP, GTP and 8 µM UTP, 2 µl of 800 Ci/mmol 10 mCi/ml 12.5 µM UTP α-^32^P (1 µM, 10-20 µCi final)(PerkinElmer), and 30 units T7 RNA polymerase in a 25 µl reaction. Next, 1 µl Turbo DNase (2 U) (Thermo-Fisher) was added to the reactions for 10 min at 37°C and then 1 µl of 250 mM of EDTA and 250 mM EGTA was added to the reaction. The probes were purified using a G25 sephadex (GE Life Sciences) spin-column. To detect 18S rRNA, 1.7 µg of 18S rRNA deoxy-oligonucleotide antisense probe (see **Supplementary File 1**) was phosphorylated using 2 µl of 6000 Ci/mmol, 150 mCi/ml, 25 µM ATP γ-^32^P (2.5 µM final, 25 to 100 µCi)(PerkinElmer) and 40 units of T4 Polynucleotide Kinase (New England Biolabs) in a 20 µl reaction incubated at 37°C for 40 minutes. The probe was then purified with G25 Sephadex column.

For anti-sense Nluc and FFluc probes, 2.5-7.5 x 10^6^ total cpm was added to the blot that had been pre-hybridized for 45 min at 68°C in 8 ml of ULTRAhyb hybridization buffer (Invitrogen). The blot was then incubated with probe at 68°C overnight, washed two times sequentially with 2 ml each of 2x SSC, 0.1% SDS, and then two more times with 0.1x SSC, 0.1% SDS at 68°C for 15 minutes each wash. For 18S rRNA probes, 5-6 x 10^6^ cpm was added to the blot that had been pre-hybridized with 8 ml ULTRAhyb-Oligo hybridization buffer (Invitrogen) at 42°C. The blot was incubated with probe overnight and then washed twice with 25 ml 2x SSC containing 0.5% SDS for 30 minutes each wash at 42°C. Blots were then exposed to phosphor screens and visualized using a Typhoon FLA phosphorimager (GE Life Sciences) and analyzed using ImageQuant TL software (GE Life Sciences). Background signal was subtracted using the “Rolling Ball” method in ImageQuant.

Nluc and FFluc levels were measured in phosphorimager units (PIU). For analysis of steady state reporter mRNA levels, fold change was determined in the same manner as described above for the reporter activity measurements, first normalizing Nluc signal to the corresponding FFluc signal in that sample, and then calculating the fold change relative to the negative control effector/condition. In tethered function assays, MS2-EGFP served as the negative control for normalization of the effectors. In the full-length Pum experiment, the RNA-binding defective mutant, Pum mut R7, served as the negative control effector.

For experiments measuring effect of endogenous Pum and corepressors on reporter mRNA levels, Northern data was analyzed in two ways. First, the values of the Pum regulated Nluc 3xPRE were divided by the Nluc ΔPRE reporter to normalize Pum specific activity to global effects on gene expression. From this data, the fold change was calculated relative to non-targeting control (NTC) negative control RNAi. In the second approach, the RRR of Nluc 3xPRE reporter mRNA was normalized to the RRR of Nluc ΔPRE reporter mRNA within each RNAi condition. From these RRR values, the fold change in Nluc 3xPRE mRNA was calculated relative to NTC.

To measure RNA decay rates, Nluc signal was normalized to stable 18S rRNA signal for each sample to adjust for potential variation in loading and transfer of RNA in each lane over the time courses. The fraction of reporter mRNA remaining at each time point was plotted relative to time in minutes after Actinomycin D addition. Half-lives were calculated using non-linear regression analysis with curve fitting to first order exponential decay. Mean mRNA half-lives and 95% credible intervals are reported for each experiment.

High resolution Northern blotting was performed to analyze Nluc 2xMS2 pA and HSL reporter mRNAs. First, 3 µg of total RNA was heated at 70°C with 20 pmol of antisense Nluc cleavage oligo RA 296 (see Supplementary Table S1) in a 30 µl reaction containing 200 mM KCl and 1 mM EDTA. In control reactions that remove the poly(A) tail (the A_0_ control) 1.5 µg of oligo deoxythymidine (dT) was included. Reactions were then cooled at room temperature for 20 minutes. Next, 5 Units of RNase H (New England Biolabs) in 20 mM Tris-HCl pH 8.0, 28 mM MgCl_2_, and 48 units of RNasin (Promega) were added and reactions were incubated at 37°C for 1 hour. Next, EDTA (final 30 mM) was added and reactions were incubated for 15 minutes at 37°C. The RNA was then purified with Clean and Concentrator–25 kit (Zymo). Next, 1.2 µg of purified RNA was combined equal volume (15 µl) of RNA loading buffer (88% formamide, 0.025% Bromophenol blue, 0.025% Xylene cyanol, 10 mM EDTA, and 0.025% SDS), heated for 10 minutes at 75° C. Samples were then electrophoretically separated on a 5% Poly-acylamide, 1 x Tris-Borate-EDTA (TBE), 8 M Urea gel (BioRad) that had been pre-run at 20-25 mA, 200 Volts, in 1x TBE buffer (89 mM Tris, 89 mM Boric acid and 2 mM EDTA). Next, the RNA was transferred onto Immobilon-Ny+ Membrane (Millipore) for 45 minutes in 0.5 x TBE Buffer at 60 V at 4°C using a Trans-Blot Cell (BioRad). The blot was crosslinked with 120 J/cm^2^ UV (λ=254) and probed with a radioactive, antisense 2x MS2 RNA probe (see **Supplementary File 1** for primers).

### Reverse Transcription and Quantitative Polymerase Chain Reaction

All parameters for RT-qPCR, including primer sequences, amplification efficiencies, and amplicon sizes, are reported according to MIQE guidelines (59) in **Supplementary File 2**. Data and statistics for RT-qPCR are reported in **Supplementary Table 1**. Reverse transcription was performed using GoScript (Promega) following the manufacturer’s instructions. Purified RNA (1.6-4 µg) and 500 ng of random hexamers were combined and heated in 10 µl volume of RNase free water at 70 °C for 5 minutes, followed by cooling on ice for 5 minutes. Next, GoScript Buffer (1x final), dNTPs (0.5 mM final), MgCl_2_ (2 mM final), and 20 units of RNase inhibitor, and 160 units GoScript reverse transcriptase were combined in a 20 µl reaction that was then incubated at room temperature for 5 minutes, 42 °C for 45 minutes, and 70 °C for 15 minutes. As a negative control for each primer set, mock “no RT” reactions were performed using identical conditions except that the reverse transcriptase was omitted. Next, qPCR was performed using GoTaq qPCR Master Mix (Promega) with the equivalent of either 50 ng or 100 ng (specified in **Supplementary File 2**) of input RNA and 0.1 µM of each primer per reaction in 50 µl final volume. In addition, no template reactions were also performed wherein cDNA was omitted, so as to assess potential false positive signal for each primer set.

The following cycling parameters were performed using a CFX96 Real-Time PCR System thermocycler (BioRad): 1) 95°C for 3 minutes, 2) 95°C for 10 seconds, 3) 65°C (for Not1, Not2, Not3, Caf40 Dcp1, and Dcp2 reactions) or 62°C (for Pop2, Ccr4, Not10, and Not11 reactions) for 30 seconds, 4) 72°C for 40 seconds, 5) repeat steps 2-4 for 40 cycles. Melt curve was generated with range 65-95°C at increments of 0.5°C. Data were analyzed using Pfaffl method (60), normalizing to the Rpl32 mRNA, with fold change calculated relative to the NTC negative control as previously described (36, 61).

## RESULTS

### Pumilio accelerates mRNA degradation

To investigate the mechanism of Pum-mediated repression, we utilized luciferase reporter gene assays in the *Drosophila* cell line, d.mel-2. We previously demonstrated that d.mel-2 cells express a limiting amount of endogenous Pum, and RNAi-mediated depletion of Pum specifically increased protein expression from a luciferase reporter mRNA bearing three PRE sequences in a minimal 3’UTR (25,36,44). We also reported that over-expression of full-length Pum protein (Figure 1A) further repressed expression of the PRE-containing reporter in a dosage dependent manner. The effect is highly specific, as mutation of the PRE sequence or the RNA recognition amino acids of the 7th repeat in the Pum RBD (Figure 1A, mut R7) alleviated RNA-binding and repression (25,36,44).

Here we report that endogenous and over-expressed Pum specifically accelerate degradation of a PRE-containing Nano-luciferase reporter mRNA. First, we analyzed the decay rate of reporters with or without PRE sequences (Figure 1B, Nluc 3xPRE versus Nluc ΔPRE) using a transcription shut-off approach and Northern blot detection (58). The PREs substantially decreased the Nluc 3xPRE mRNA level and reduced its half-life by 7.3-fold relative to Nluc ΔPRE, with a P(diff)>0.999 (i.e. posterior probability that the two half-life values are different)(Figure 1C and 1D), indicating that Pum recognition of the mRNA caused its degradation.

We note that the statistical analyses in this report utilized a Bayesian approach that incorporates a multilevel model to assess contributions to error across multiple technical, biological, and experimental replicates. Data reported in figures include mean values and 95% credible intervals, with posterior probabilities and number and type of replicates reported in the figure legends and **Supplementary Table 1**. Following previous work on Pum proteins (62), we assess differences based on 95% credible intervals and P(sig), the posterior probability of a difference of at least 1.3-fold in the direction indicated. High values of P(sig) indicate high confidence in a biologically meaningful effect being present (P(sig)≥0.95 are indicated by a ‘**’ in the figures, whereas differences that are significant but do not meet this 1.3-fold threshold are marked with a ‘*’). We also use a converse measure, P(insig), which we define as the posterior probability that a change is no larger than 1.3-fold ((P(insig)≥0.95 are indicated by a ‘x’ in the figures). We note that a large value of P(insig) is far more informative than a large p-value would be in a frequentist statistical test; whereas the latter only represents a failure to reject the null hypothesis, and thus has limited inferential value, a large value of P(insig) represents true confidence that an effect is small.

We next demonstrated that PRE-mediated mRNA decay is dependent on Pum. RNAi depletion of Pum, verified by RT-qPCR in Figure 1E, increased the half-life of the Nluc 3xPRE mRNA by 5.3-fold (P(diff)=0.99) relative to non-targeting control RNAi (Figure 1F and 1G). Because the Pop2 deadenylase subunit of the CNOT complex is implicated in Pum-mediated repression, we tested the effect of Pop2 depletion, confirmed in Figure 1E, and observed a 3.5-fold increase in Nluc 3xPRE mRNA half-life (P(diff)>0.999) (Figure 1F and G). This result indicates that Pum-PRE-mediated mRNA decay occurs through the deadenylation-mediated pathway.

Next, we measured the effect of over-expressed Pum on mRNA decay. We previously showed that Pum expression specifically repressed PRE-containing reporter mRNAs, whereas an RNA-binding defective mutant, Pum mut R7, did not (25,36,44). Consistent with those observations, expression of wild type Pum, but not mut R7, repressed protein expression from the Nluc 3xPRE mRNA by more than 4-fold (P(sig)>0.999) in a dual luciferase reporter assay (Figure 1H and 1I) and accelerated mRNA decay (Figure 1J and 1K), reducing the Nluc 3xPRE mRNA half-life by 2.3-fold (P(diff)>0.999). Expression of the V5 epitope-tagged Pum effector proteins was confirmed by western blot of equal mass of cell extracts (Figure 1I). Together, these results indicate that Pum-PRE mediated repression accelerates mRNA degradation and that Pop2 deadenylase is important for decay of PRE-bearing mRNAs.

### N-terminal Pumilio Repression Domains cause mRNA decay

We previously showed that the N-terminus of Pum confers its major repressive activity, mediated by three Repression Domains (Figure 1A, RD1, RD2, and RD3) (36). These domains can repress in a tethered function assay, wherein they are fused to the RNA-binding domain of the MS2 phage coat protein and directed to the 3’UTR of a reporter mRNA bearing tandem MS2 binding sites, Nluc 2xMS2 (Figure 2A) (36,63–65). These assays include a co-transfected control Firefly luciferase gene (FFluc) for normalization of transfection efficiency in each sample. In agreement with our previous observations (36, 44), tethering the Pum N-terminus or each RD reduced reporter protein expression (Figure 2B, Nluc 2xMS2), whereas the negative control MS2-EGFP fusion did not. Under these conditions, Pum N and RD1 repressed by 2-fold relative to MS2-EGFP, while RD2 and RD3 repressed by 1.4-fold and 1.5-fold, respectively (all Pum effectors had P(sig)>0.999). As a positive control, the decapping enzyme subunit Dcp1, fused to MS2, elicited 3.9-fold repression (P(sig)>0.999)(Figure 2B), consistent with the previously reported effect of tethering decapping factors (53,66–70). Importantly, the observed repression was dependent on binding of effector proteins to the reporter, because they had little or no effect on a reporter that lacks the MS2 binding sites, Nluc ΔMS2 (each Pum effector had P(insig)>0.999)(Figure 2B). We also compared the repressive activity of each effector between the two reporters, demonstrating that Pum N repressed by 3.3-fold (P(sig)>0.999), RD1 by 3.1-fold (P(sig)>0.999), RD2 by 2-fold (P(sig)=0.99), RD3 by 2.5-fold (P(sig)>0.999), and the Dcp1 control repressed by 6.5-fold (P(sig)>0.999)(Supplementary Figure S1A). The expression of the V5 epitope-tagged effector proteins was confirmed by western blotting of equal mass of cellular lysates (Figure 2C). These results support the independent repressive activity of each Pum RD.

**Figure 2.**
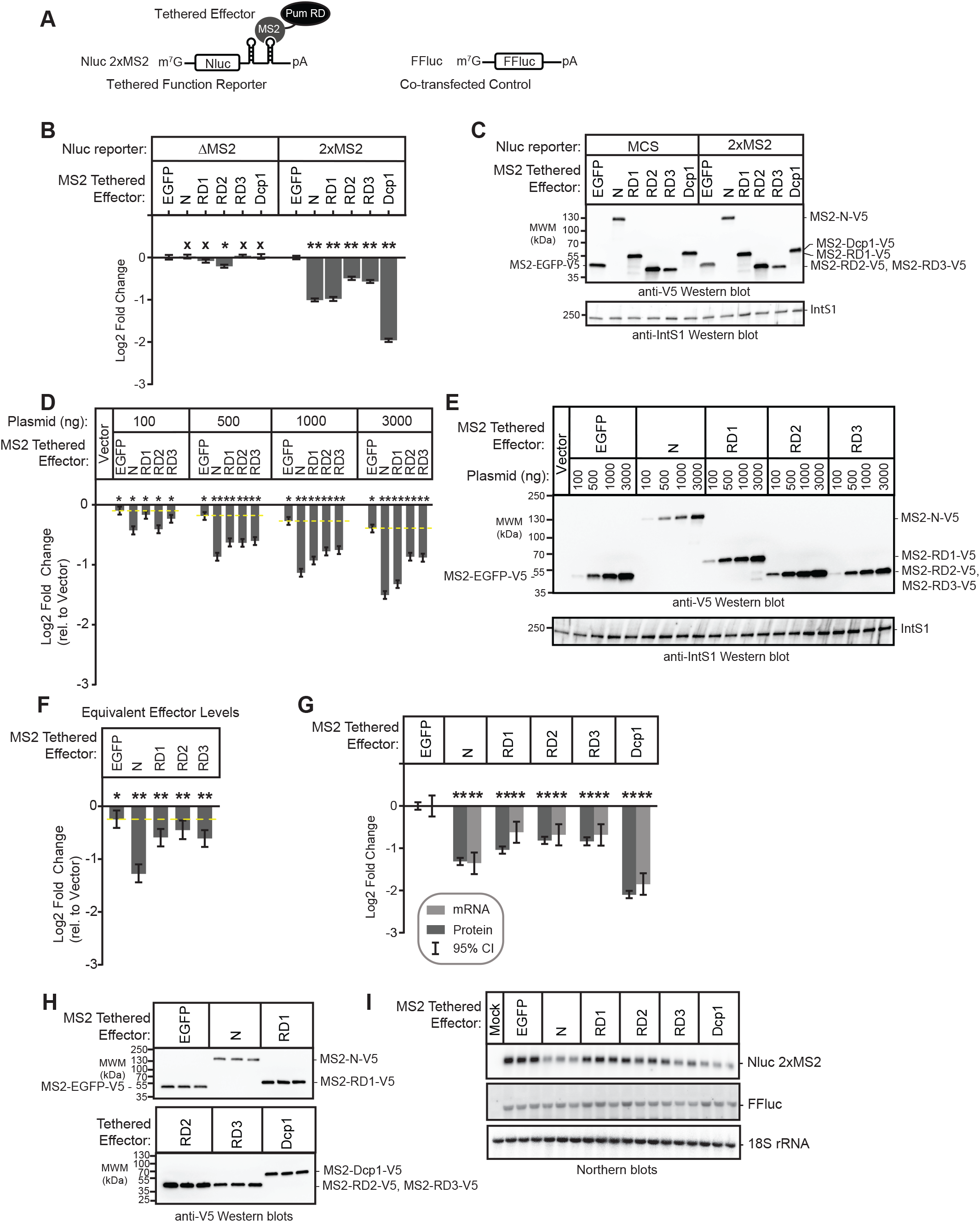
The Pumilio N-terminal Repression Domains repress mRNA and protein expression. (A) Diagram of tethered function Nano-luciferase reporter mRNA (Nluc 2xMS2) and co-transfected control, Figure Firefly luciferase (FFluc). Nluc 2xMS2 bears two copies of the MS2 stem loop RNA structure in its 3’UTR, which is bound by the sequence-specific MS2 RNA-binding protein. By expressing Pum or other effectors as a fusion to MS2 RNA-binding protein, the impact of the effector on reporter protein and mRNA levels can be measured. Diagram is not to scale. (B) The repression activity of Pum N-terminus and individual repression domains (RD1, RD2, RD3) was measured using the tethered function dual luciferase assay using the Nluc 2xMS2 reporter or an equivalent reporter wherein the MS2 binding sites are deleted, Nluc ΔMS2. Mean log_2_ fold change values in normalized reporter activity, measured relative to the negative control, MS2-EGFP, are plotted with 95% credible intervals, from 3 experiments with 4 technical replicates each. Tethered decapping enzyme subunit, Dcp1, serves as a positive control that strongly represses the reporter when tethered. Data and statistics are reported in Supplementary Table S1. Comparison of the activity of each Pum effector on the Nluc 2xMS2 reporter relative to Nluc ΔMS2 is shown in Supplementary Figure S1A. For significance calling, a ‘*’ denotes a posterior probability >0.95 that the difference relative to the negative control is in the indicated direction. The ‘**’ indicates a posterior probability of >0.95 that the indicated difference is at least 1.3-fold. An ‘x’ marks a posterior probability >0.95 that the indicated difference is no more than 1.3-fold in either direction. (C) Western blot of the V5-epitope tagged MS2 fusion effector proteins used in panel B from a representative experimental replicate. Equivalent mass of protein from each sample was probed with anti-V5 antibody, followed by the Integrator Subunit 1 protein (IntS1) to assess equal loading of lanes. (D) The relationship of repression activity to level of each MS2 tethered effector was measured using the tethered function assay by titrating the amount of transfected effector expression plasmid, as indicated at the top. Total mass of transfected plasmid was maintained across all conditions by supplementing with empty expression plasmid vector, pIZ. In this experiment, repression of reporter activity was calculated relative to the control condition containing only the empty expression vector. Data from three experiments with four technical replicates each, are plotted along with 95% credible intervals. The yellow dashed line demarks the repression activity beyond that of the negative control effector, MS2-EGFP, for each amount of transfected effector plasmid. The relationship of effector protein level, measured by quantitative western blot, and repression activity is reported in Supplementary Figure S1B. Data and statistics are reported in Supplementary Table S1. (E) Western blot of the V5-epitope tagged MS2 fusion effector proteins used in panel D from a representative experimental replicate. Equivalent mass of protein from each sample was probed with anti-V5 antibody, followed by IntS1 as a loading control. (F) The repression activity for equivalent expression level of each MS2 tethered effector protein, determined by tethered function assays and quantitative western blotting as shown in Supplementary Figure S1B. Fold change was calculated relative to empty vector. Mean log_2_-fold change values are plotted with 95% credible intervals from 3 experiments with 4 technical replicates each. The yellow dashed line demarks the repressive activity beyond the negative control effector, MS2-EGFP. Data and statistics are reported in Supplementary Table S1. (G) The effect of each MS2 fusion effector protein on Nluc 2xMS2 reporter protein and mRNA level was determined by dual luciferase assay (reporter Protein, dark grey bars) and Northern blotting (reporter mRNA, light grey bars). Log_2_ fold change of Nluc 2xMS2 levels, normalized to internal control FFluc, for each effector were calculated relative to negative control MS2-EGFP for three biological replicates. Mean log_2_ fold change and 95% credible intervals are reported in the graph. Data and statistics are reported in Supplementary Table S1. (H) Expression of V5-tagged MS2 fusion effector proteins from three biological replicates (panel G) was confirmed by western blotting of an equal mass of protein cell extract for each sample. (I) Northern blot detection of tethered function reporter Nluc 2xMS2, internal control FFluc, and loading control 18S rRNA for three biological replicate samples for each tethered effector protein. Each lane of the gel contains 5 µg of total RNA. This data was used to determine fold change in reporter mRNA level shown in panel G. The Mock sample contained total cellular RNA from untransfected cells and demonstrates specificity of the reporter probes.

To further characterize the activities of the N-terminal repression domains, we analyzed the relationship of effector protein dosage to repression of Nluc 2xMS2 reporter activity. To do so, the mass of transfected effector expression plasmid (MS2-Pum N, MS2-RD1, MS2-RD2, MS2-RD3, or MS2-EGFP) was titrated over a 30-fold range and repression was measured relative to cells transfected with empty expression vector, pIZ (Figure 2D and 2E). To maintain identical transfection conditions, the total mass of transfected DNA was balanced across samples with empty expression vector. Pum N and RDs exhibited repression activities proportional to the mass of expression plasmid, and substantially above the equivalent amount of negative control MS2-EGFP (Figure 2D).

We then examined the relationship of effector protein level to repression by performing quantitative western blotting on equal mass of cell extract from these samples (Figure 2E). We observed a log-linear relationship wherein increased Pum effector protein level caused a proportional decrease in reporter expression (Supplementary Figure S1B). In contrast, MS2-EGFP was far less effective. The only substantial deviation from a log-linear relationship of effector amount to repressive activity occurs in the case of MS2-RD1, for which higher effector protein levels appear to be somewhat disproportionately more effective (Supplementary Figure S1B). Importantly, repression activity by each Pum effector was observed to be consistent and proportional across a broad range of plasmid or protein levels; even a 30-fold reduction in effector did not eliminate repression by Pum effectors (Figure 2D and Supplementary Figure S1B).

To directly compare repression activities, the fold change relative to empty vector was determined for equivalent amounts of expressed effector proteins. This analysis shows that per unit of effector protein, the order of efficacy is: Pum N > RD3 = RD1 = RD2 > EGFP (Figure 2F). The posterior probability of the difference of Pum N is greater than the RDs is P(sig)>0.999 and the activities among the three RDs are not distinguishable from each other (P(insig)≥0.93), whereas their activities are consistently greater than the EGFP negative control (P(sig)≥0.97) (**Supplementary Table 1**). We conclude that the tethered function assay provides a robust and specific means of assaying Pum RD activity. Minor fluctuations in effector level do not result in loss of activity, nor do they alter our qualitative conclusions regarding the effects of the various constructs.

To measure the impact of each effector on Nluc 2xMS2 mRNA, we performed Northern blot analysis. The Pum N-terminus and RDs reduced the Nluc reporter mRNA level with magnitudes corresponding to their effects on reporter protein level (P(sig)≥0.97)(Figure 2G-2I), whereas the internal control Firefly luciferase mRNA was not affected by the tethered effectors (Figure 2I). Northern blot of the 18S ribosomal RNA served as an internal control for gel loading and blotting. These results indicate that the N-terminal Pum RDs promote mRNA decay.

### The putative Pumilio cap-binding motif is not required for repression

We interrogated a model wherein Pum was hypothesized to repress translation via a 5’ cap-binding motif located in its N-terminus (11, 48). This motif does not correspond to the three RDs, but instead resides within a conserved region (previously designated PCMb)(36). First, we introduced a mutation in the Pum N-terminus, W783G (Figure 1A), which was reported to disrupt cap binding (48) and compared its repressive activity (Figure 3A) and expression (Figure 3B) to wild type in the tethered function assay. Both wild type and mutant Pum N-terminus repressed the reporter with equivalent effectiveness (e.g. 3000 ng of either wild type or mutant W783G Pum N repressed by 2.9-fold with P(sig)=1.0, relative to EGFP negative control) and in a dose dependent manner (Figure 3A). By direct comparison of the repressive activities of wild type Pum N to the mutant W783G, we see that there is no significant difference in their repressive activities (P(insig)>0.999), for each transfected amount.

**Figure 3.**
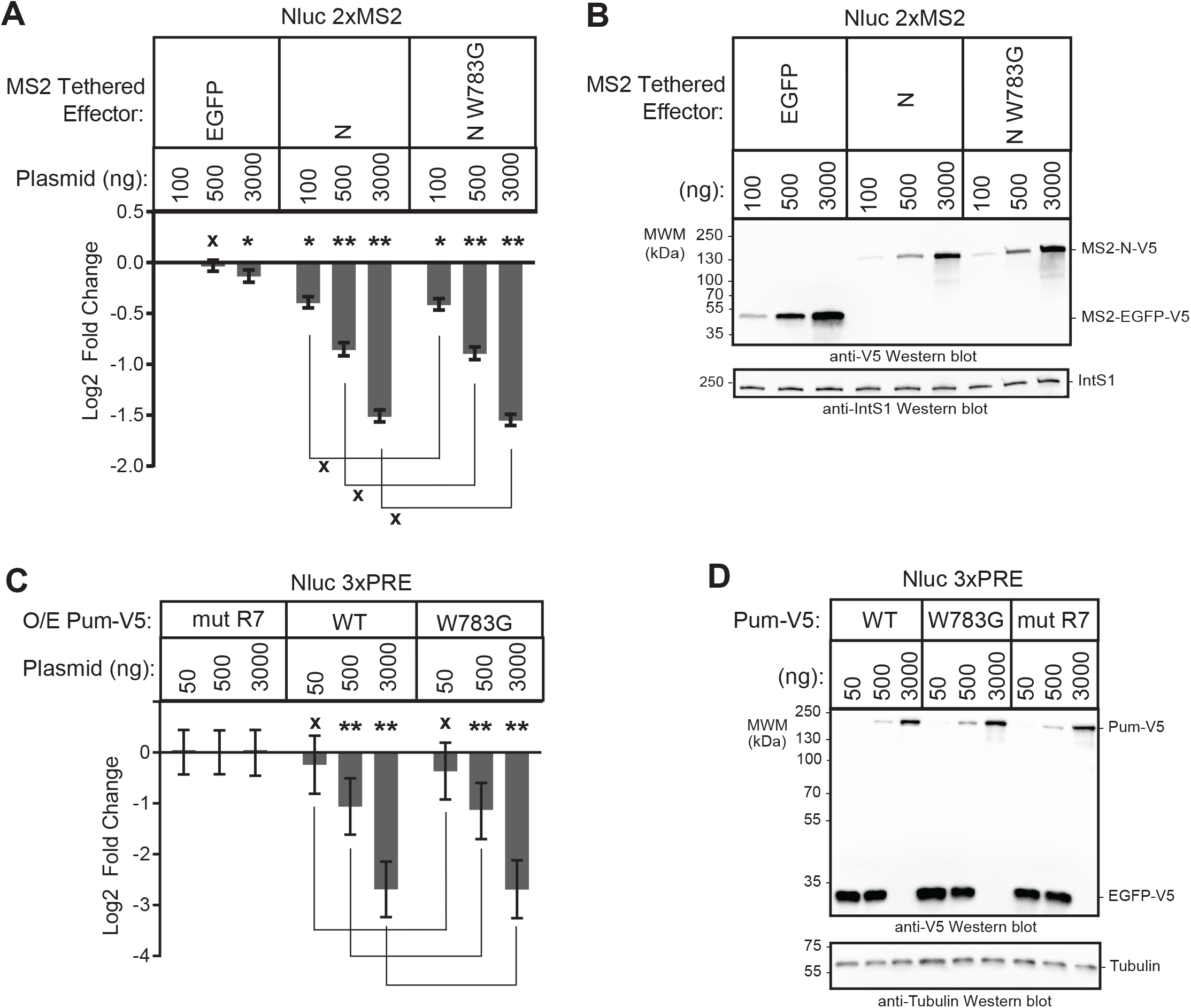
The putative Pumilio cap-binding motif is not required for repression. (A) Repression activity was measured for three amounts of transfected wild type or cap-binding mutant (W873G) Pum N-terminus via the tethered function dual luciferase assay using the Nluc 2xMS2 pA reporter. Repression activity was calculated relative to the MS2-EGFP negative control at the lowest transfected amount (100 ng). Empty expression vector, pIZ, was used to balance the total mass of transfected plasmids in samples with 100 and 500 ng of MS2 effector plasmid. Mean log_2_ fold change and 95% credible intervals for three experimental replicates with four technical replicates each are reported in the graph. Data and statistics are reported in Supplementary Table S1. For significance calling, a ‘*’ denotes a posterior probability >0.95 that the difference relative to the negative control is in the indicated direction. The ‘**’ indicates a posterior probability of >0.95 that the indicated difference is at least 1.3-fold. An ‘x’ marks a posterior probability >0.95 that the indicated difference is no more than 1.3-fold in either direction. (B) Western blot of the V5-epitope tagged MS2 fusion effector proteins used in panel A from a representative experimental replicate. Equivalent mass of protein from each sample was probed with either anti-V5 antibody or anti-IntS1 as a loading control. (C) Repression activity was measured for three amounts of transfected wild type or cap-binding mutant (W873G) full length Pum via dual luciferase assay using the Nluc 3xPRE pA reporter. The fold change values were calculated relative to the equivalent amount of transfected RNA-binding defective mutant Pum (mut R7) negative control. V5-tagged EGFP plasmid served to balance the total mass of transfected plasmids in samples with 50 and 500 ng of Pum effector plasmid. Mean log_2_ fold change and 95% credible intervals for three experimental replicates with four technical replicates each are reported in the graph. Data and statistics are reported in Supplementary Table S1. (D) Western blot of the V5-epitope tagged Pum effector and EGFP balancer proteins used in panel C from a representative experimental replicate. Equivalent mass of protein from each sample was probed with anti-V5 antibody and, to assess equal loading of lanes, anti-Tubulin antibody.

We also examined the effect of the W783G mutation on repression of a PRE-containing reporter by over-expressed full length Pum (Figure 3C and 3D); no significant reduction in the ability of the mutant Pum to repress was observed. For example, 3000 ng of either wild type Pum or mutant Pum W783G repressed by 6.4-fold with P(sig)=1.0, relative to the mutR7 negative control. Based on this data, and our previous observation that PCMb was neither necessary nor sufficient for repression (36), we conclude that - in these experimental conditions - the proposed 5’ cap binding motif does not contribute to Pum-mediated repression.

### CNOT complex components are important for Pumilio Repression Domain activity

We sought to identify co-repressors necessary for repression by the Pum RDs. The CNOT deadenylase complex plays a crucial role in the initiation of mRNA decay (39) and is important for mRNA decay by Pum-PRE (Figure 1F-G), thus we evaluated its role in repression by Pum N-terminus and individual RDs. The *Drosophila* CNOT complex contains 8 subunits and the Pop2 subunit is thought to be the major deadenylase (Figure 4A)(40); therefore, we first performed RNAi using two different double stranded RNAs (dsRNAs) to deplete Pop2. DsRNA1 targets the open reading frame of Pop2 mRNA, whereas the dsRNA2 targets its 5’UTR. Both dsRNAs depleted Pop2 from the d.mel-2 cells relative to non-targeting control dsRNA (NTC), as confirmed by RT-qPCR (Figure 4B) and Western blotting (Figure 4C). Pop2 depletion was effective and reproducible in each condition using either dsRNA, though depletion by dsRNA1 was more robust (Figure 4B).

**Figure 4.**
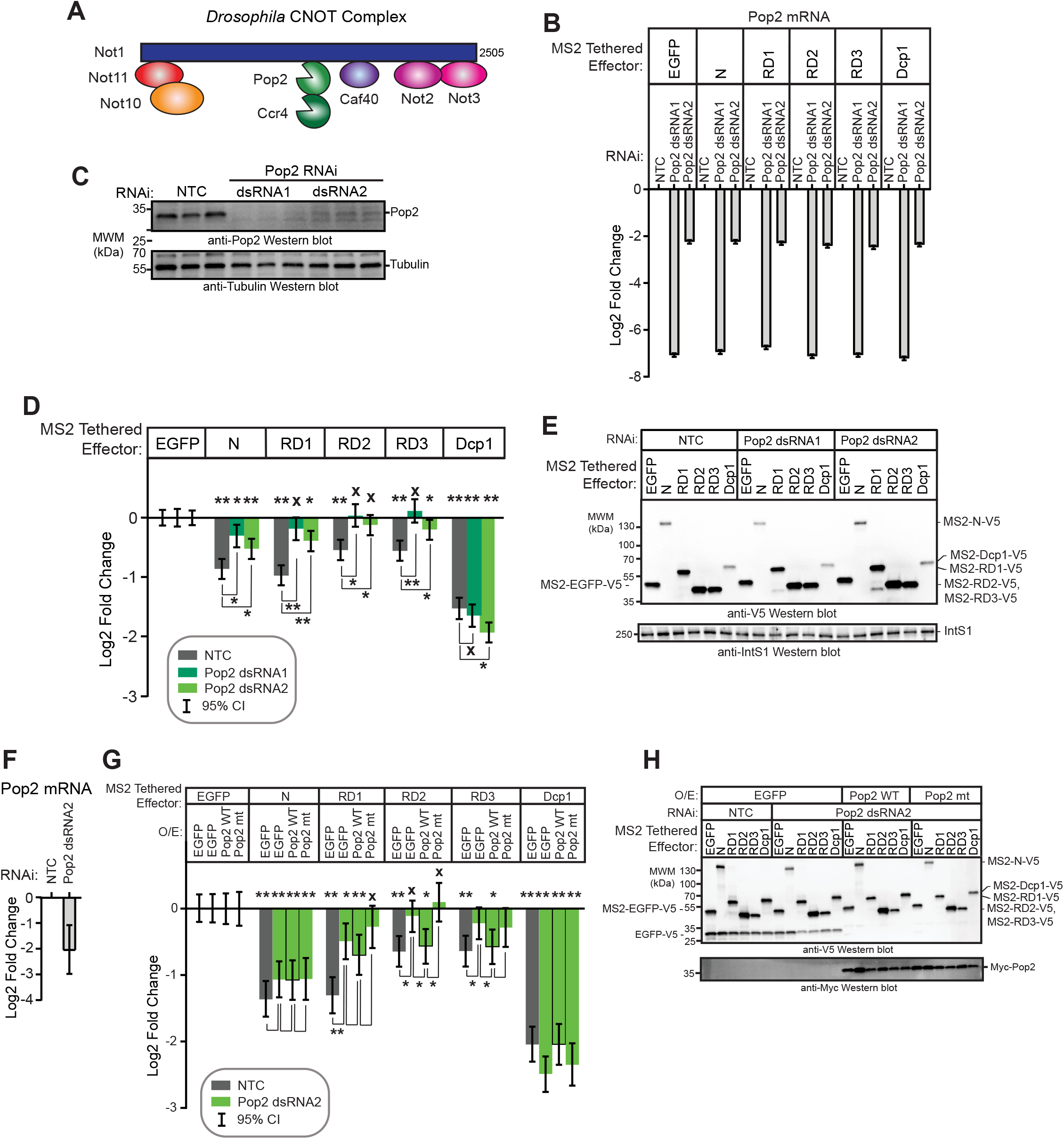
The Pop2 deadenylase is required for Pum RD activity. (A) Diagram of the *Drosophila melanogaster* Ccr4-Not complex, containing 8 subunits. Adapted from Temme et al. (71). (B) The efficiency of Pop2 mRNA depletion after 3 days of treatment with either of two double stranded RNAs (dsRNA1 and dsRNA2) was measured using RT-qPCR. The dsRNA1 targets the Pop2 coding sequence, whereas dsRNA2 targets the Pop2 mRNA 5’UTR. Fold changes were calculated relative to non-targeting control RNAi (NTC) for the indicated experimental conditions. Mean log_2_ fold change and 95% credible intervals for a representative experimental replicate with three technical replicates for each measurement are reported in the graph. Data and statistics are reported in Supplementary Table S1. (C) Western blot confirming RNAi-mediated depletion of Pop2 deadenylase induced by treatment of d.mel-2 cells (3 biological replicates) with two different double-stranded RNAs in comparison to non-targeting control dsRNA (NTC). Equivalent mass of cellular extract was loaded for each sample. Anti-tubulin western blot serves as a loading control. (D) The effect of Pop2 depletion on repression by Pum N-terminus and individual RDs was measured via tethered function dual luciferase assay. Repression by each effector was calculated relative to the corresponding negative control effector MS2-EGFP within each RNAi condition. Non-targeting control (NTC) RNAi serves as a negative control for RNAi. Tethered decapping enzyme subunit, Dcp1, serves as a positive control. Mean log_2_ fold change and 95% credible intervals for three experimental replicates with four technical replicates each are reported in the graph. Data and statistics are reported in Supplementary Table S1. For significance calling, a ‘*’ denotes a posterior probability >0.95 that the difference relative to the negative control is in the indicated direction. The ‘**’ indicates a posterior probability of >0.95 that the indicated difference is at least 1.3-fold. An ‘x’ marks a posterior probability >0.95 that the indicated difference is no more than 1.3-fold in either direction. (E) Western blot of the V5-epitope tagged MS2 fusion effector proteins used in panel D from a representative experimental replicate. Equivalent mass of protein from each sample was probed with anti-V5 antibody, followed by western blot of IntS1 as a loading control. (F) The efficiency of Pop2 mRNA depletion after 5 days of Pop2 dsRNA2 treatment was measured using RT-qPCR. Fold changes were calculated relative to non-targeting control (NTC). Mean log_2_ fold change and 95% credible intervals for three biological replicates with three technical replicates each are reported in the graph. Data and statistics are reported in Supplementary Table S1. (G) The ability of wild type Pop2 (WT) or active site mutant Pop2 (mt) to rescue repression by Pum N-terminus and RDs was measured via tethered function dual luciferase assay. Endogenous Pop2 was depleted by treating cells with dsRNA2. NTC dsRNA serve as a control. The effect of Pop2 expression was compared to EGFP control. Mean log_2_ fold change and 95% credible intervals from 3-6 experimental replicates with four technical replicates each are reported in the graph. Data and statistics are reported in Supplementary Table S1. (H) Western blot of V5-tagged tethered effectors and myc-tagged Pop2, mutant Pop2, or negative control V5-tagged EGFP from a representative experimental replicate in panel G. Equivalent mass of cellular extract was loaded for each sample.

We then measured the effect of Pop2 depletion on repression by tethered Pum N-terminus and RDs. It is important to note that in all reporter gene assays that incorporate RNAi, the repressive activity of the effector was measured relative to the negative control effector, MS2-EGFP, within the same RNAi condition, as described in the Methods. In this manner, the specific effect of RNAi-mediated depletion on the Pum effector is determined. We observed that Pop2 depletion substantially reduced but did not eliminate repression by the Pum N-terminus, decreasing from 1.8-fold in the NTC condition (P(sig)=1) to 1.2-fold by Pop2 dsRNA1 (P(sig)=0.21) (Figure 4D). Pop2 depletion eliminated repressive activity of RD2 and RD3 (P(sig)=0) relative to MS2-EGFP), and greatly reduced RD1 activity from 2-fold in the NTC condition (P(sig)=1) to 1.1-fold with Pop2 dsRNA1 (P(sig)=0.03)(Figure 4D). As anticipated based on the ability of Dcp1 to interact with the mRNA decapping enzyme Dcp2, Pop2 depletion did not alleviate repression by tethered decapping enzyme subunit Dcp1, which maintained repressive activity in all conditions (P(sig)=1.0)(Figure 4D). Effector expression was verified in each condition (Figure 4E). Based on this data, we conclude that Pop2 is important for repressive activity of the Pum N-terminus and RDs.

To investigate whether the deadenylase activity of Pop2 is involved in Pum RD mediated repression, we tested the ability of wild type or mutant Pop2 to rescue repression. To do so, cells were treated with Pop2 dsRNA2 and then were transfected with plasmid expressing RNAi resistant cDNAs encoding myc-tagged wild type or catalytically inactive Pop2 mutant (D53A and E55A) (40), or V5-tagged EGFP as a control. Depletion of Pop2 mRNA was verified by RT-qPCR (Figure 4F). As before, depletion of Pop2 reduced repression by the Pum effectors in the control EGFP expressing conditions (Figure 4F, compare Pop2 dsRNA2 to NTC). Importantly, expression of wild type Pop2, but not the active site mutant Pop2, increased repression activity of RD2 and RD3 in cells treated with Pop2 dsRNA2 (Figure 4G). RD1 activity also increased, but the change did not meet our significance threshold. No effect of Pop2 expression on the activity of Pum N was evident. Protein expression of all effectors and the wild type and mutant Pop2 proteins was confirmed by western blot analysis of equivalent mass of cell extracts in Figure 4H. These observations indicate that the deadenylation activity of Pop2 is important for RD activity.

Ccr4 is the second deadenylase present in the CNOT complex (Figure 4A) (41, 71). We next evaluated the involvement of Ccr4 and observed that substantial depletion of its mRNA (Supplementary Figure S2A) did not affect repression by the Pum N-terminus or RDs (P(insig)≥0.91)(Supplementary Figure S2B and S2C). This observation suggests that Ccr4 is not crucial, whereas Pop2 has an important role in Pum repression. That said, the potential limitation of RNAi is notable, wherein residual low levels of Ccr4 might be sufficient to support Pum activity.

We also examined the role of non-catalytic CNOT subunits in Pum RD-mediated repression, starting with Not1, which is the central scaffold of the complex (Figure 4A) (71). RNAi depletion of Not1 was confirmed by RT-qPCR (Figure 5A) and western blotting (Figure 5B). We observed that knockdown of Not1 abrogated repression by RD2 (decreasing from 1.5-fold (P(sig)=0.88 to 1.1-fold (P(sig)=0.04) and RD3 (decreasing from 1.5-fold (P(sig)=0.92) to 1.1-fold (P(sig)=0.11)(Figure 5C). Moreover, Not1 depletion substantially reduced repression by RD1 (decreasing from 2-fold (P(sig)=1) to 1.4-fold (P(sig)=0.68) and the entire N-terminus (decreasing from 2.2-fold (P(sig)=1) to 1.6-fold (P(sig)=0.99)(Figure 5C).

**Figure 5.**
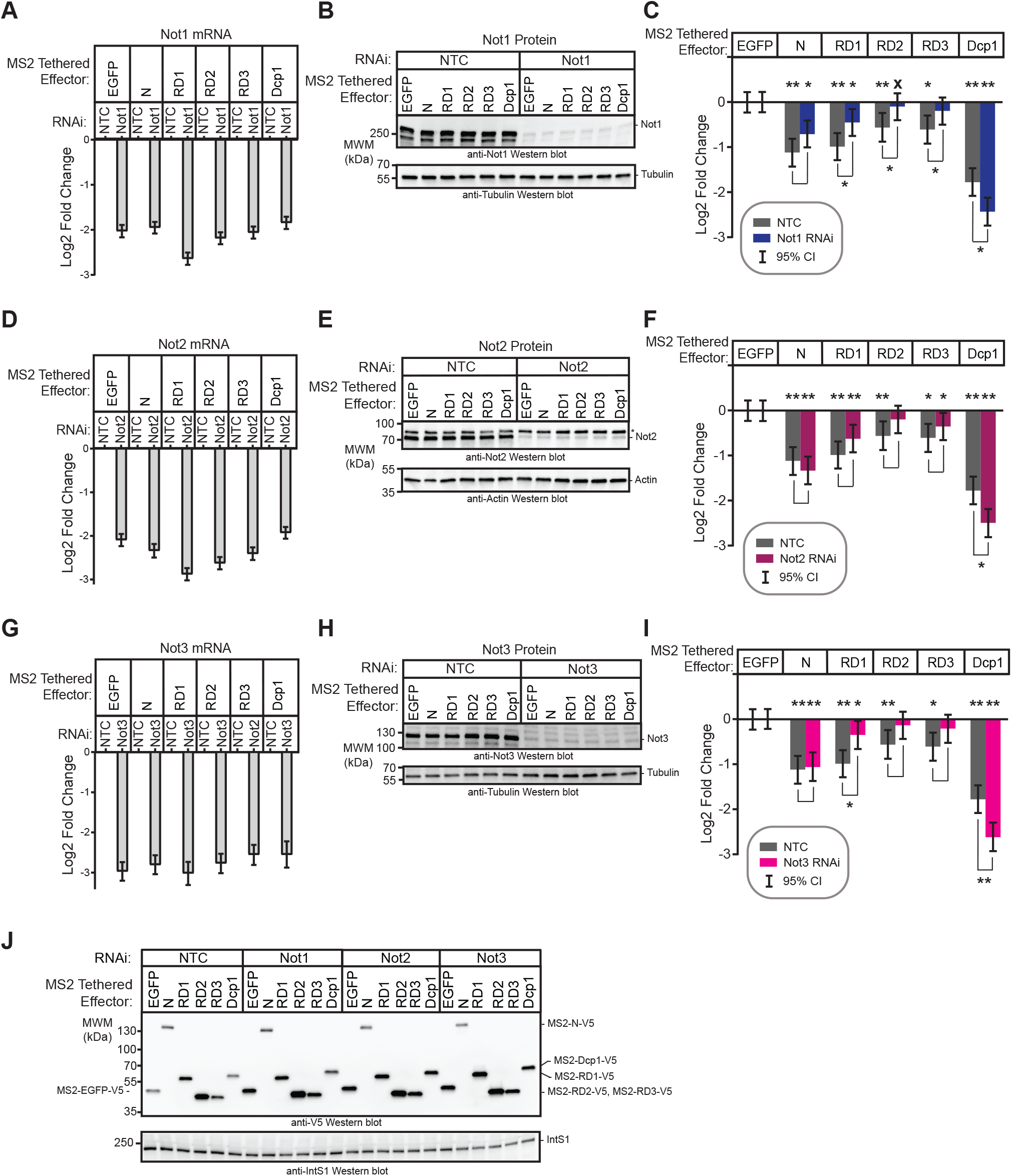
CNOT components are involved in Pum RD mediated repression. (A) The efficiency of RNAi-mediated depletion of Not1 mRNA after 3 days of dsRNA treatment was measured using RT-qPCR. Fold changes were calculated relative to non-targeting control (NTC). Mean log_2_ fold change and 95% credible intervals are reported in the graph for one representative experiment with three technical replicates of each measurement. Data and statistics are reported in Supplementary Table S1. (B) Western blot with anti-Not1 antibody confirms depletion of endogenous Not1 protein from a representative experiment. Equivalent mass of cellular extract was loaded for each sample. Anti-tubulin western blot serves as a loading control. (C) Tethered function dual luciferase assays measured the effect of Not1 depletion on the repression activity of Pum N-terminus and RDs. Non-targeting control (NTC) serves as negative control for comparison. Activity of each effector was determined relative to the corresponding negative control effector MS2-EGFP within each RNAi condition. Mean log_2_ fold change and 95% credible intervals are reported in the graph for three experimental replicates with four technical replicates each. Data and statistics are reported in Supplementary Table S1. For significance calling, a ‘*’ denotes a posterior probability >0.95 that the difference relative to the negative control is in the indicated direction. The ‘**’ indicates a posterior probability of >0.95 that the indicated difference is at least 1.3-fold. An ‘x’ marks a posterior probability >0.95 that the indicated difference is no more than 1.3-fold in either direction. (D) The efficiency of Not2 mRNA depletion after 3 days of dsRNA treatment was measured using RT-qPCR. Fold changes were calculated relative to non-targeting control (NTC). Mean log_2_ fold change and 95% credible intervals are reported in the graph for one representative experiment with three technical replicates of each measurement. Data and statistics are reported in Supplementary Table S1. (E) Western blot confirms RNAi-mediated depletion of endogenous Not2 from a representative experiment using an anti-Not2 antibody. Equivalent mass of cellular extract was loaded for each sample. Anti-actin western blot serves as a loading control. The * designates a protein that cross-reacts with the Not2 antibody. (F) Tethered function dual luciferase assays measured the effect of Not2 depletion on the repression activity of Pum N-terminus and RDs. Non-targeting control (NTC) serves as negative control for comparison. Activity of each effector was determined relative to the corresponding negative control effector MS2-EGFP within each RNAi condition. Mean log_2_ fold change and 95% credible intervals are reported in the graph for three experimental replicates with four technical replicates each. Data and statistics are reported in Supplementary Table S1. (G) The efficiency of Not3 mRNA depletion after 3 days of dsRNA treatment was measured using RT-qPCR. Fold changes were calculated relative to non-targeting control (NTC). Mean log_2_ fold change and 95% credible intervals are reported in the graph for one representative experiment with three technical replicates of each measurement. Data and statistics are reported in Supplementary Table S1. (H) Western blot confirms RNAi depletion of endogenous Not3 protein from a representative experiment using an anti-Not3 antibody. Equivalent mass of cellular extract was loaded for each sample. Anti-tubulin western blot serves as a loading control. (I) Tethered function assays measured the effect of Not3 depletion on the repression activity of Pum N-terminus and RDs. Non-targeting control (NTC) serves as negative control for comparison. Activity of each effector was determined relative to the corresponding negative control effecto MS2-EGFP within each RNAi condition. Mean log_2_ fold change and 95% credible intervals are reported in the graph. Data and statistics are reported in Supplementary Table S1. (J) Western blot of the V5-epitope tagged MS2 fusion effector proteins used in panels C, F and I from a representative experiment. Equivalent mass of protein from each sample was probed with anti-V5 antibody and IntS1 loading control.

We observed that RNAi depletion of Not1 also reduced the level of its protein partner Pop2 (Supplementary Figure S3A and S3B), consistent with a previous report (40). Therefore, the effect of RNAi of Not1 on Pum activity could be the result of diminished Pop2. To test this idea, we attempted to rescue the effect of Not1 depletion by expressing Pop2 from a transfected plasmid. First, we titrated Pop2 expression vector to approximate the level of endogenous Pop2 protein (Supplementary Figure S3B). We then performed tethered function assays. Again, depletion of Not1 reduced (for Pum N and RD1) or eliminated (for RD2 and RD3) repression activity (Supplementary Figure S3C). Expression of exogenous Pop2 did not rescue the loss of repression by the Pum N-terminus or RDs caused by Not1 depletion. Based on these observations, we conclude that Not1 is important for Pum RD activity, and its depletion mimics the effect of Pop2 depletion.

The other CNOT subunits form modules that interact with Not1, as depicted in Figure 4A (71). We observed that depletion of Not2 (Figures 5D-F) and Not3 (Figures 5G-I) also reduced Pum RD-mediated repression. Overall, depletion of Not2 or Not3 uniformly reduced the activity of Pum RDs comparably to depletion of Not1, although in some instances the effect did not meet our statistical significance threshold (Figure 5F and I). Pum effector expression in each RNAi condition was confirmed by western blotting (Figure 5J). We also tested the effects of depletion of Caf40, Not10, and Not11 (Supplementary Figure S2D-S2I), none of which alleviated the repressive activity of the Pum N-terminus or individual RDs (P(insig)≥0.88). We note that Caf40 knockdown did affect RD2, wherein its repressive activity was enhanced, an observation that is currently not understood. Overall, these results indicate that certain CNOT subunits are important for the repression activity of Pum RDs (i.e. Pop2, Not1, 2 and 3) whereas others appear to be dispensable (i.e. Ccr4, Caf40, Not10 and 11).

### CNOT is required for efficient Pumilio-mediated mRNA decay

Given the importance of CNOT for Pum RD activity, we further examined its involvement in Pum-mediated mRNA degradation. To do so, we depleted Pop2 by RNAi and measured decay of the Nluc 3xPRE reporter in response to over-expressed Pum. In the control RNAi condition (NTC), over-expressed wild type Pum accelerated decay of the reporter mRNA, reducing its half-life by 2.7-fold relative to the negative control, RNA-binding defective mut R7 (Figure 6A and 6B). Depletion of Pop2 stabilized the reporter RNA in the presence of WT (by an estimated 13-fold, P(diff)>0.999) or mut R7 (by an estimated 69-fold, P(diff)>0.999)(Figure 6A and 6B). This result demonstrates that Pop2 is required for efficient Pum-mediated mRNA decay, consistent with the observation that Pop2 is important for PRE-mediated decay in Figure 1F and 1G. Using the same strategy, we analyzed the involvement of Not1 and observed that depletion of Not1 also led to impairment of Pum-mediated mRNA decay (Figure 6C and 6D), stabilizing the reporter mRNA by 9.2-fold (P(diff)>0.999) in the wild type Pum condition and by more than an estimated 200-fold (P(diff)>0.999) in the mut R7 condition. This data indicates that the CNOT complex plays a crucial role in mechanism by which Pum accelerates mRNA decay.

### Pumilio N-terminal Repression Domains bind to the CNOT complex

The observation that the repressive activity of Pum N-terminal RDs require CNOT components suggested a model wherein they act to recruit the CNOT complex to target mRNAs. We used a co-immunoprecipitation assay to assess a possible physical interaction between flag-tagged Pum N-terminus or RBD with endogenous CNOT. Flag-tagged GST served as a negative control. Several positive controls were also utilized including the RNA-binding protein Nanos, which was previously shown to directly contact Not1 (45), and the stoichiometric CNOT complex subunits Not2 and Not3. We observed that the Pum N-terminus associates with Not1 (Figure 7A), similar to Nanos, and that this interaction is resistant to treatment with RNase (Figure 7A and 7B), indicating that the association is not bridged by RNA. As expected, Not1 robustly co-purified with Not2 and Not3. Interestingly, while the Pum RBD was previously shown to interact with Pop2 (42, 44), we did not detect co-immunoprecipitation with Not1 under these conditions, perhaps reflecting differences in protein-protein contacts or affinities. Together, these observations indicate that the Pum N terminus associates with CNOT.

We then investigated the potential for direct interaction between the Pum RDs and the CNOT complex by performing in vitro protein interaction assays using bead-bound purified Pum RD1, RD2, RD3, or RBD proteins, which were purified as fusions to maltose binding protein (MBP) and the StrepII affinity tag. Bead-bound MBP-StrepII served as a negative control. The eight subunit human CNOT complex was purified as described by Raisch et al. (Figure 7C and 7D**, Input**)(57). We observed that each Pum RD interacted with the intact CNOT complex, including both deadenylases. The RBD similarly interacted with the CNOT complex, consistent with its reported interaction with Pop2 (42, 44). This data indicates that each Pum repression domain can directly bind to the CNOT deadenylase complex. It is noteworthy that the CNOT complex is highly conserved throughout eukarya (39), as are Pumilio orthologs (1), and we previously reported that human Pumilio proteins are active in *Drosophila* cells (36,44,72). The observation that *Drosophila* Pum RDs bind to human CNOT accentuates the probable conservation of the repressive mechanism.

### The poly(A) tail is necessary for maximal activity of the Pum N-terminus

As a complementary approach to analyze the role of deadenylation in the mechanism of repression by the Pum N-terminus, we tested whether the poly(A) tail is necessary. To do so, we compared Pum repression of the polyadenylated Nluc 2xMS2 reporter to that of a non-adenylated reporter bearing a 3’ Histone Stem Loop (HSL) processing signal (Figure 8A) (73). First, we confirmed that each reporter generated the correct 3’ end product by cleaving them with RNase H and an antisense deoxyoligonucleotide that is complementary to the Nluc coding region (Figure 8A). In addition, deadenylated 3’ end RNA fragments were generated for each reporter by adding oligo deoxythymidine (oligo dT) to the indicated samples (Figure 8B). The resulting 3’ end fragments were detected by high resolution Northern blotting, thereby verifying that the Nluc 2xMS2 pA reporter produced a distribution of poly(A) lengths spanning up to 200 adenosines appended to the 228 nucleotide 3’ end product, whereas the Nluc 2xMS2 HSL reporter mRNA produced the expected non-adenylated 210 nucleotide product (Figure 8B).

We then compared the ability of the Pum N-terminus to repress the adenylated and non-adenylated reporters. Repression by Pum N was significantly reduced from 3.4-fold (P(sig)=1) on the polyadenylated reporter to 1.4-fold (P(sig)=0.69) on the HSL reporter (Figure 8C and 8D). Two conclusions can be drawn from this result. First, the reduction in activity emphasizes the importance of the poly(A) tail in maximal repression by the Pum N-terminus. Second, the residual repressive activity of Pum N on the non-adenylated mRNA indicates that it wields an additional poly(A) independent repressive mechanism. We also analyzed the individual Pum RDs and observed that RD1 behaved similarly to the N-terminus, while RD2 and RD3 exhibited comparable repressive activity on the two reporters (Figure 8C and 8D), indicating that the RDs contribute to poly(A) independent repression. The tethered Dcp1 control repressed both reporters (Figure 8C and 8D), consistent with its ability to promote 5′ decapping of the transcripts.

We postulated that the poly(A) independent repression activity of the N-terminus and RDs may be mediated via CNOT recruitment. This hypothesis is based on previous studies that reported deadenylation-independent repression by the CNOT complex (74, 75). Moreover, the CNOT complex interacts with translational repressors and mRNA decay enzymes, including the decapping enzyme complex (71,74,76). We therefore analyzed the role of CNOT in poly(A) independent repression by the Pum N-terminus by measuring the impact of Not1 depletion. As in Figure 5, RNAi of Not1 significantly reduced repression by Pum N on the polyadenylated reporter mRNA, but had only a modest effect on the poly(A)-independent repression of the HSL reporter (Figure 8E and 8F). In contrast, when we examined the effect of Not1 knockdown on the individual RDs, we found that their ability to repress the HSL reporter was abrogated (Figure 8G and 8H). Indeed, depletion of Not1 eliminated RD1 repression of Nluc 2xMS2 HSL from 1.3-fold (P(sig)=0.53) to 1.0-fold (P(sig)=0), RD2 repression from 1.6-fold (P(sig)=1.0) to 1.0-fold (P(sig)=0), and RD3 repression from 1.2-fold (P(sig)=0.49) to 1.0-fold (P(sig)=0). Our interpretation of these observations is that the Pum RDs utilize the CNOT complex to cause both poly(A) dependent and independent repression.

### The Pum N-terminus utilizes an additional CNOT-independent repression activity

The residual repressive activity of the Pum N-terminus on the non-adenylated HSL reporter (Figure 8), and when CNOT components are depleted (Figure 4, 5, and 8), indicates that an additional mechanism contributes to repression. We further assessed the CNOT independent activity by simultaneously depleting both Not1 and Pop2. As before, depletion of each co-repressor individually decreased Pum N activity (Figure 9A). Simultaneous depletion of Not1 and Pop2 further reduced repression (Figure 9A), from 2.2-fold (P(sig)=1) in the NTC condition to 1.4-fold (P(sig)=0.74) in the Pop2+Not1 RNAi condition. These results emphasize the importance of CNOT in poly(A) dependent repression, but also further support the residual CNOT independent repression activity.

To further understand CNOT independent repression by the Pum N-terminus, we measured its effect on the levels of Nluc 2xMS2 pA mRNA when Pop2 or Not1 are depleted. As shown in Figure 9C-E, depletion of either CNOT component diminished Pum N mediated reduction of reporter mRNA levels, decreasing from 5.2-fold (P(sig)>0.999) repression of mRNA level in the NTC control condition to 1.7-fold (P(sig)>0.98) or 2.3-fold (P(sig)>0.999), when Pop2 or Not1 were depleted, respectively. Again, Pum N retained residual ability to reduce reporter mRNA levels. These observations are consistent with the results in Figures 1 and 6, wherein CNOT depletion stabilized the Pum-repressed mRNA, but did not completely eliminate decay.

**Figure 6.**
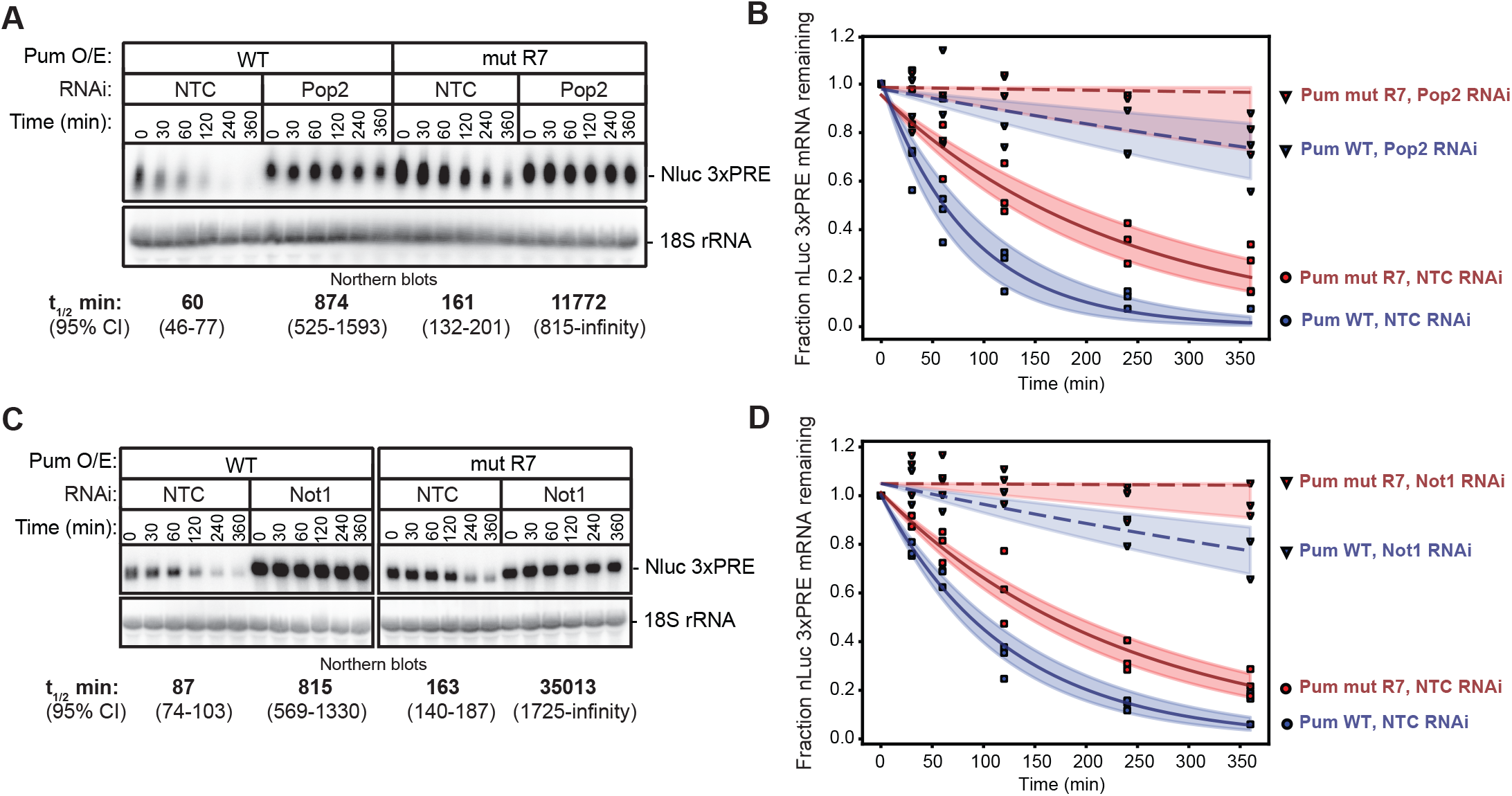
Pum-mediated mRNA decay requires Not1 and Pop2. (A) The effect of RNAi-mediated depletion of Pop2 by Pop2 dsRNA1 on the mRNA decay rate of Nluc 3xPRE reporter mRNA was measured in response to over-expressed wild type Pum (Pum WT) or the RNA-binding defective mutant Pum (Pum mut R7) following inhibition of transcription with ActD. Cells treated with NTC dsRNA served as negative control. The Nluc 3xPRE was detected by Northern blot along with 18S ribosomal RNA (rRNA), as a loading control. Each lane of the gel contains 10 µg of total RNA. The mRNA half-lives and 95% credible intervals measured in each condition are shown below the respective blots, and were calculated from three experimental replicates. Data and statistics are reported in Supplementary Table S1. (B) The fraction of Nluc 3xPRE mRNA remaining, normalized to 18S rRNA, is plotted relative to time (minutes) after inhibition of transcription. Data points for each of 3 experimental replicates are plotted. First order exponential decay trend lines, calculated using non-linear regression analysis, are plotted for each effector (Pum and mut R7) in each RNAi condition (NTC, dark solid lines, and Pop2, dashed solid lines), along with the 95% credible interval for each trend line (shaded areas: orange for mutR7 and blue for Pum WT). (C) The effect of RNAi-mediated depletion of Not1 on the mRNA decay rate of Nluc 3xPRE reporter mRNA was measured in response to over-expressed wild type Pum (Pum WT) or the RNA-binding defective mutant Pum (Pum mut R7) following inhibition of transcription with ActD. Cells treated with NTC dsRNA served as negative control. The Nluc 3xPRE was detected by Northern blot along with 18S ribosomal RNA (rRNA), as a loading control. Each lane of the gel contains 10 µg of total RNA. The mRNA half-lives and 95% credible intervals measured in each condition are shown below the respective blots, and were calculated from three experimental replicates. Data and statistics are reported in Supplementary Table S1. (D) The fraction of Nluc 3xPRE mRNA remaining, normalized to 18S rRNA, is plotted relative to time (minutes) after inhibition of transcription. Data points for each of 3 experimental replicates are plotted. First order exponential decay trend lines, calculated using non-linear regression analysis, are plotted for each effector (Pum and mut R7) in each RNAi condition (NTC, dark solid lines, and Not1, dashed solid lines), along with the 95% credible interval for each trend line (shaded areas: orange for mut R7 and blue for Pum WT).

### Decapping enzyme participates in repression by the Pum N-terminus

We postulated that decapping may play a role in the residual repression activity of the Pum N-terminus. Removal of the 5’ cap of mRNAs plays a key role in the destruction of mRNAs through the 5’ decay pathway (38), and decapping is catalyzed by the enzyme Dcp2, which forms a complex with Dcp1 (76). We therefore inhibited decapping and examined its role in repression by the Pum N-terminus. To do so, Dcp2 was depleted via RNAi (as verified by RT-qPCR in Figure 10A) while simultaneously over-expressing an RNAi-resistant, catalytically inactive, dominant negative Dcp2 mutant (E361Q). As previously established (45,52,53), this combined approach was necessary to effectively abrogate decapping-mediated mRNA decay.

Inhibition of decapping diminished repression of reporter protein expression by the Pum N-terminus from 2.2-fold (P(sig)=1) to 1.7-fold (P(sig)=1)(Figure 10B) and also alleviated the ability of the N-terminus to reduce reporter Nluc 2xMS2 pA mRNA level from 2.7-fold (P(sig)=1) to 1.2-fold (P(sig)=0.4)(Figure 10B and 10C), indicating that decapping participates in the repression mechanism of Pum N-terminus. Supporting the efficacy of the approach, inhibition of decapping stabilized the Nluc reporter mRNA and Firefly luciferase internal control mRNA (Figure 10C). Further corroboration is provided by the observation that mRNA degradation by tethered Dcp1 was prevented by inhibition of Dcp2 (Figure 10B and 10C). Interestingly, tethered Dcp1 retained the ability to repress reporter translation, which likely reflects its association with translational inhibitory factors (Figure 10B) (76). Based on this collective data, we conclude that decapping contributes to the repression activity of the Pum N-terminus.

### Multiple mechanisms contribute to Pumilio-mediated repression

Having identified multiple co-repressors – Not1, Pop2, Dcp2 (this study) and pAbp (44) – that are important for the individual activities of multiple Pum repression domains (N-terminal RDs and the C-terminal RBD), we investigated their contributions to regulation by full-length, endogenous Pum. To do so, Pum repression was measured using the Nluc 3xPRE reporter and compared to the unregulated reporter Nluc ΔPRE, which lacks PRE sequences. As before, FFluc mRNA served as an internal control. Not1, Pop2, Dcp2, or pAbp were each depleted by RNAi, as confirmed by western blot (Figure 11A) or RT-qPCR (Figure 11B), and then reporter protein (Figure 11C and 11D) and mRNA levels (Figure 11E) were measured by dual luciferase assay and Northern blot, respectively. As before, the RNAi depletion of Dcp2 was accompanied by over-expression of mutant Dcp2 E361Q.

The resulting data was analyzed in two ways (Figure 11C and 11D). First, the Relative Response Ratio (RRR) was calculated for each sample by dividing the Nluc reporter signal by the internal control FFluc signal. This approach normalizes potential sample-to-sample variation in transfection efficiency, and also responds to global effects on gene expression. Next, to specifically measure PRE-dependent regulation in each RNAi condition, the RRR value of the Nluc 3xPRE reporter was divided by that of the RRR unregulated Nluc ΔPRE reporter. Then, the fold change values for each RNAi condition, reported in Figure 11C, were calculated relative to the non-targeting control (NTC) RNAi condition. These results measure the PRE specific effect of depletion of the Pum co-repressors on reporter protein and mRNA levels. RNAi depletion of Pum served as a positive control (Figure 11A) and alleviated PRE-dependent repression, resulting in increased reporter protein (2.8-fold, P(sig)>0.99) and mRNA levels (2.4 fold, P(sig)>0.99) (Figure 11C). Depletion of Not1 or Pop2 alleviated PRE-dependent repression (2.1-fold, P(sig)>0.99 and 2.0-fold, P(sig)>0.99, respectively) and stabilized the reporter mRNA (1.6-fold, P(sig)=0.98 and 1.9-fold, P(sig)=0.86, respectively) (Figure 11C). Inhibition of Dcp2 more modestly increased reporter protein expression (1.4-fold, P(sig)=0.99) and stabilized the PRE-containing reporter mRNA (1.5-fold, P(sig)=0.83) (Figure 11C), as did depletion of pAbp (1.4-fold increase in PRE reporter protein, P(sig)=0.99; 1.3-fold increase in PRE reporter mRNA, P(sig)=0.44), reflecting their respective contributions to repression by the Pum N-terminus and RBD.

Consistent with their global roles in basal mRNA decay, depletion of CNOT, and to a lesser degree Dcp2, affected FFluc levels, as observed by Northern blot (Figure 11E). Hence, we also analyzed PRE-dependent regulation by omitting the internal control FFluc from the calculations. In this case, to specifically measure PRE-dependent regulation, the level of the Nluc 3xPRE reporter was divided by the unregulated Nluc ΔPRE reporter for each RNAi condition, and the fold change values were then calculated relative to the negative control (NTC) RNAi condition. The resulting PRE-dependent fold change values are reported in Figure 11D. Again, depletion of Pum (2.2-fold increase, P(sig)>0.99), Not1 (2.0-fold increase, P(sig)>0.99), or Pop2 (2.1-fold increase, P(sig)>0.99) resulted a substantial loss in PRE-dependent repression. Depletion of pAbp caused a 1.4-fold increase (P(sig)=0.91) and Dcp2 inhibition had a minor effect (1.1-1.2 fold increase, though without normalization to transfection efficiency, the variation limited our ability to draw a firm conclusion). Taken together, our results demonstrate the multiple co-repressors contribute to Pum-mediated repression of target mRNA and protein levels in *Drosophila* cells.

## DISCUSSION

The results of this study provide new insights into the molecular mechanism by which Pum represses gene expression. Previous work correlated Pum repression with reduction in mRNA levels, and we now show that Pum uses multiple repression domains to accelerate mRNA decay via the CNOT deadenylase and decapping complexes. This information enhances our understanding of the biological roles and impact of Pum on the transcriptome.

The model that emerges from our findings is that Pum utilizes four domains that use the CNOT deadenylase complex to repress target mRNAs (Figure 11F). Multiple lines of evidence directly link CNOT to Pum-mediated repression. First, Pum requires CNOT subunits to repress protein expression and accelerate RNA decay, specifically the Not1, Not2, Not3, and Pop2 subunits (Figures 4 and 5). Second, the catalytic activity of Pop2, the major deadenylase enzyme in *Drosophila*, is required to rescue Pum RD-mediated repression in the Pop2 RNAi condition (Figure 4). Third, the poly(A) tail is necessary for maximal repression by Pum (Figure 8). Fourth, CNOT co-immunoprecipitates with the Pum N-terminus in an RNase resistant manner (Figure 7). Fifth, the Pum RDs and RBD directly bind to the CNOT complex (Figure 7). In addition, the C-terminal RBD of Pum was previously shown to interact with Pop2 and promote deadenylation (42, 44). Moreover, Pum has been linked to deadenylation of target mRNAs during *Drosophila* development (8, 43).

**Figure 7.**
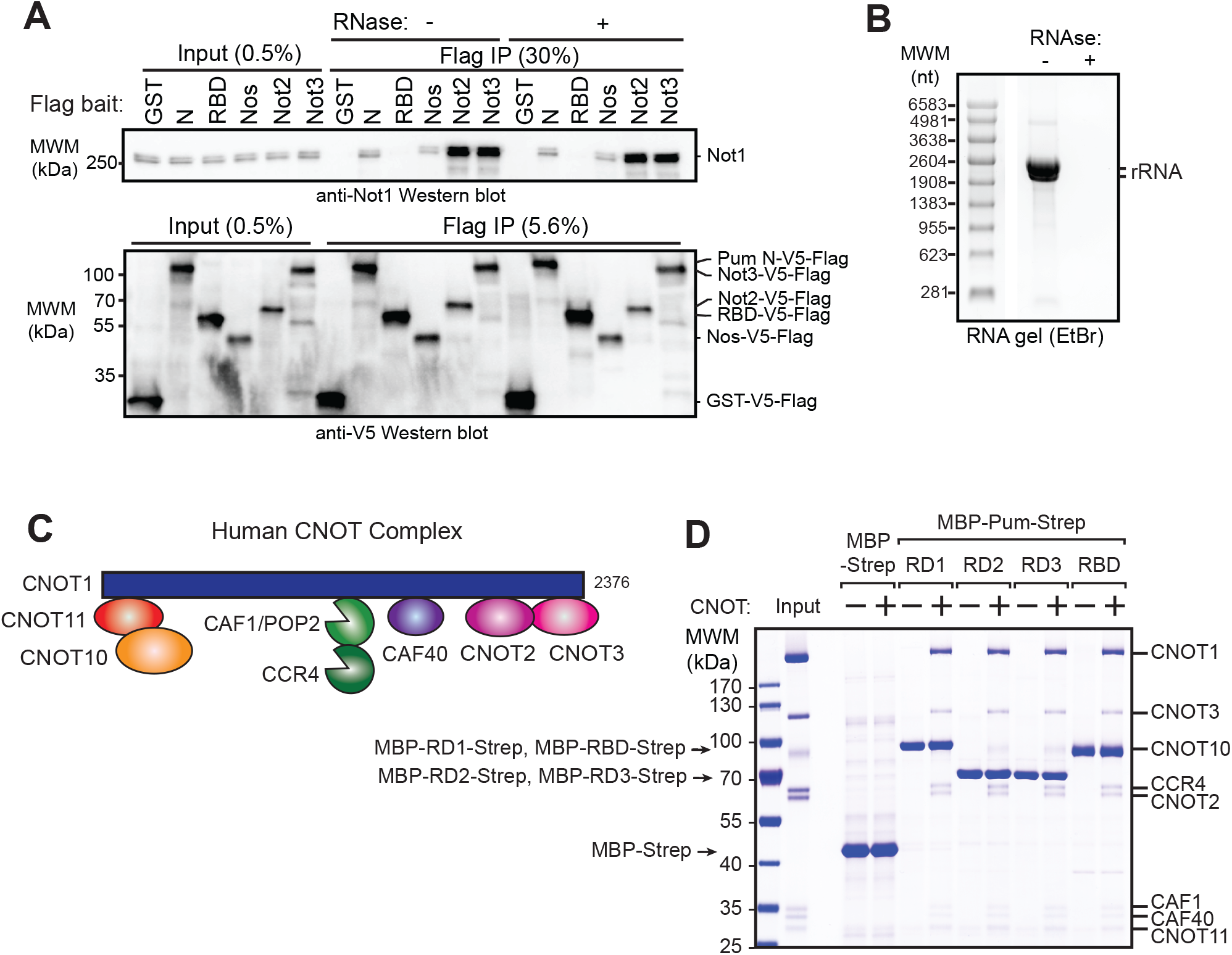
Pumilio N-terminal Repression Domains bind to the CNOT complex. (A) Not1 protein co-immunoprecipitates with the Pum N-terminus from d.mel2 cell extracts. Western blot detection of endogenous Not1 protein and Flag- and V5-tagged Pum N-terminus (N) or RNA-binding domain (RBD) in cellular extracts (Input) and anti-Flag immunoprecipitates (Flag IP) from samples treated with (+) or without (-) RNase One treatment of the cellular extracts. Flag-V5-tagged GST serves as negative control. Positive controls for Not1 interaction include Flag-V5-tagged Nanos (Nos) and core CNOT subunits Not2 and Not3. The relative percent of total Input and Flag IP for each sample is indicated above lanes. (B) Confirmation of RNA digestion by RNase One in co-immunoprecipitation experiment in panel A. Total RNA purified from the cellular extract treated with (+) or without (-) RNase One was analyzed on denaturing formaldehyde agarose gel and visualized with ethidium bromide. Ribosomal RNA is indicated in the sample without RNase treatment. Note that *Drosophila* 28S rRNA (3945 nt) is internally processed to two fragments (∼1787 nt and ∼2112 nt) whereas the 18S rRNA is ∼1995 nt (112, 113). (C) Diagram of the human Ccr4-Not complex containing 8 subunits. Note that the subunits are orthologous – compare Figure 4A and 7C – though the nomenclature differs between human and *Drosophila* as described in Temme et al, 2014 (71). (D) Pum RDs and RBD bind to the intact human CNOT complex. In vitro protein interaction “pulldown” assays were performed using recombinant, purified, streptactin bead-bound Pum domains (indicated at the top) that were fused to maltose binding protein (MBP) and the StrepII affinity tag (Strep). Bead-bound MBP-Strep serve as a negative control. Human CNOT complex (Input), purified as described by Raisch et al, 2019 (57), was incubated with the bead bound bait proteins. After extensive washing, bead bound proteins were analyzed by Coomassie blue-stained SDS-PAGE. A representative experiment of three experimental replicates is shown.

Why does Pum use multiple domains to recruit CNOT? Perhaps their activities combine to increase the avidity of Pum for CNOT, mediated by multiple contacts between Pum domains and CNOT. Indeed, the four repressive domains each directly interact with the CNOT complex (Figure 7). Mapping the precise protein-protein interactions necessary for CNOT recruitment by Pum will require detailed biochemical and structural analysis, to be pursued in future studies.

Recruitment of CNOT by RNA-binding factors has emerged as an important mechanism of post-transcriptional regulation (39,45,77–80). Utilization of CNOT by Pum orthologs has been reported in *Saccharomyces cerevisiae*, *Schizosaccharomyces pombe*, *Caenorhabditis elegans*, *Drosophila melanogaster* and mammals (11,43,44,72,81–83), and thus represents an evolutionarily conserved mechanism. Like *Drosophila* Pum, the highly conserved RBD of Pum orthologs spanning from yeast to human universally interact with Pop2 orthologs. In contrast, the N-terminal RDs are a more recent evolutionary addition, being found in Pum orthologs spanning insects through mammals (36). Based on the results of this study, we speculate that analogous regions of those Pum orthologs repress by recruiting the CNOT complex. Consistent with this idea, experimental evidence showed that the N-termini of human Pum orthologs, PUM1 and PUM2, exhibit repressive activity when directed to an mRNA (36). Future research should dissect the repressive mechanism of mammalian Pum N-termini.

It is interesting to speculate that Pum might recruit a subcomplex of CNOT, based on the observation that Pop2, Not1, Not2, and Not3 were important for Pum RD activity whereas depletion of others had little to no effect (i.e. Ccr4, Not10, Not11, Caf40)(Supplementary Figure S2). Germane to this idea, evidence in yeast and human cells revealed heterogeneity in size and composition of CNOT complexes (84–87). The requirement of specific CNOT subunits for repression by Pum orthologs was also observed in *S. cerevisiae* (82, 88). Still, interpretation of the negative results in our experiments is limited by the effectiveness of RNAi-mediated depletion, and we acknowledge that residual levels of a CNOT component may be sufficient to support activity. Moreover, our biochemical analysis indicates that Pum repression domains can bind the intact CNOT complex (Figure 7).

We also found that decapping contributes to repression by Pum, supported by several lines of evidence. First, the N-terminus retains partial repressive activity when the target mRNA lacks a poly(A) tail and therefore is not subject to deadenylation (Figure 8). In addition, the N-terminus retains repression and RNA decay activities when Not1 and/or Pop2 are depleted (Figures 1, 6, and 9). Moreover, inhibition of the decapping enzyme Dcp2 reduced repression and RNA decay activity of the Pum N terminus (Figures 10 and 11). Future analyses will be necessary to delineate the specific region(s) of Pum that modulate decapping and how it associates with decapping enzyme – either via direct protein contacts or via bridging proteins that interface with the decapping machinery (76). Interestingly, decapping appears to be a conserved mechanism of Pum repression, supported by the observations that yeast PUF proteins associate with decapping factors and promote decapping of target mRNAs (82,89,90).

**Figure 8.**
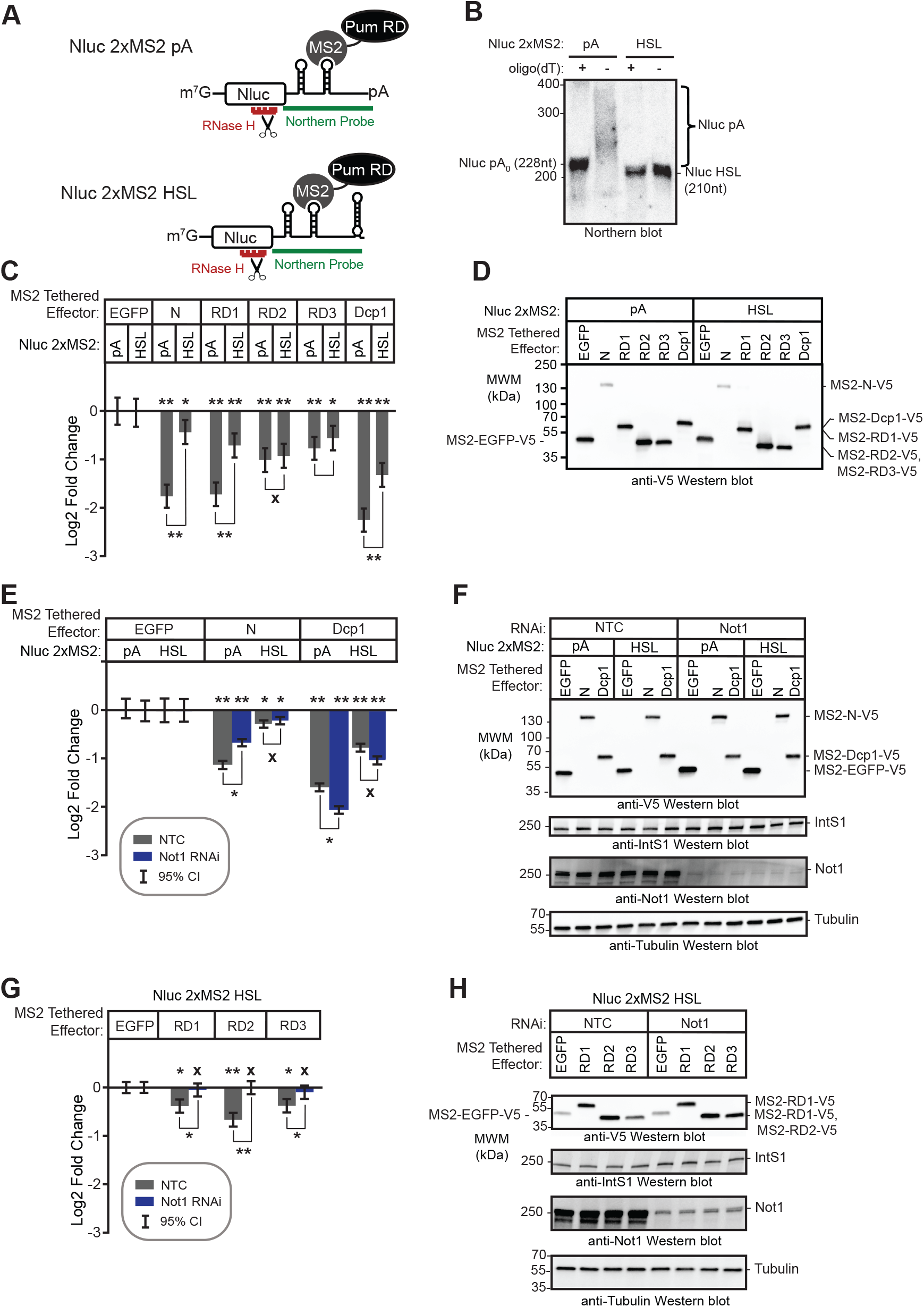
The poly(A) tail is necessary for maximal activity of the Pum N-terminus. (A) Diagram of the Nluc 2xMS2 reporters with either 3′ poly(A) tail or the Histone Stem Loop (HSL) used for tethered function assays. The reporters are identical except that the cleavage/poly-adenylation element was replaced with a Histone Stem Loop and Histone Downstream Element (HDE) in the Nluc 2xMS2 HSL reporter, which produces a non-adenylated 3′ end. The location of the probe used for Northern blotting (green) and the DNA oligonucleotide (red) used for RNase H cleavage of the mRNAs for high resolution Northern blotting are indicated. Diagram is not drawn to scale. (B) High resolution Northern blot of the Nluc 2xMS2 pA and HSL reporter mRNAs expressed in d.mel2 cells confirms proper poly-adenylation of the pA reporter and lack of poly-adenylation of the HSL reporter. Where indicated (+), RNA was treated with DNA oligonucleotide of 15 thymidines (dT) and RNase H to degrade the poly(A) tail. RNA size markers are indicated on the left. The lengths of the 3′ end reporter fragments are also indicated. (C) The repression activity of the Pum N-terminus and RDs was measured via tethered function dual luciferase assay, using the Nluc 2xMS2 poly(A) and HSL reporters. The activity of each effector was determined relative to the tethered EGFP negative control effector on the same reporter. Mean log_2_ fold change and 95% credible intervals are reported in the graph for three experimental replicates with four technical replicates each. Data and statistics are reported in Supplementary Table S1. For significance calling, a ‘*’ denotes a posterior probability >0.95 that the difference relative to the negative control is in the indicated direction. The ‘**’ indicates a posterior probability of >0.95 that the indicated difference is at least 1.3-fold. An ‘x’ marks a posterior probability >0.95 that the indicated difference is no more than 1.3-fold in either direction. (D) Western blot detection of the V5-epitope tagged MS2 fusion effector proteins used in panel C from a representative experimental replicate. Equivalent mass of protein from each sample was probed with anti-V5 antibody. (E) The effect of RNAi-mediated depletion of Not1 on repression activity of the Pum N-terminus was measured via tethered function, using the Nluc 2xMS2 poly(A) and HSL reporters. The non-targeting control (NTC) served as negative control for RNAi. The repression activity of each effector was determined relative to tethered EGFP negative control in the same RNAi condition. The mean log_2_ fold change and 95% credible intervals are graphed for three experimental replicates with four technical replicates each. Data and statistics are reported in Supplementary Table S1. (F) Western blot of the V5-epitope tagged MS2 fusion effector proteins from a representative experimental replicate from panel E. Equivalent mass of protein from each sample was probed with anti-V5 antibody or anti-IntS1 to assess equal loading of lanes. The depletion of Not1 protein was assessed using an anti-Not1 antibody, with anti-tubulin western blot serving as the loading control. (G) The effect of RNAi depletion on the repression activity of the Pum RDs was measured via tethered function dual luciferase assay, using the Nluc 2xMS2 HSL reporter. Non-targeting control (NTC) serves as negative control for comparison. Activity of each effector was determined relative to tethered EGFP negative control in the same RNAi condition. The mean log_2_ fold change and 95% credible intervals are reported in the graph for three experimental replicates with four technical replicates each. Data and statistics are reported in Supplementary Table S1. (H) Western blot of the V5-epitope tagged MS2 fusion effector proteins used in from a representative experimental replicate from pane G. An equivalent mass of protein from each sample was probed with anti-V5 antibody, and then with anti-IntS1 to assess equal loading of lanes. Depletion of the Not1 protein was assessed using an anti-Not1 antibody, with ant-tubulin western blot serving as a loading control.

**Figure 9.**
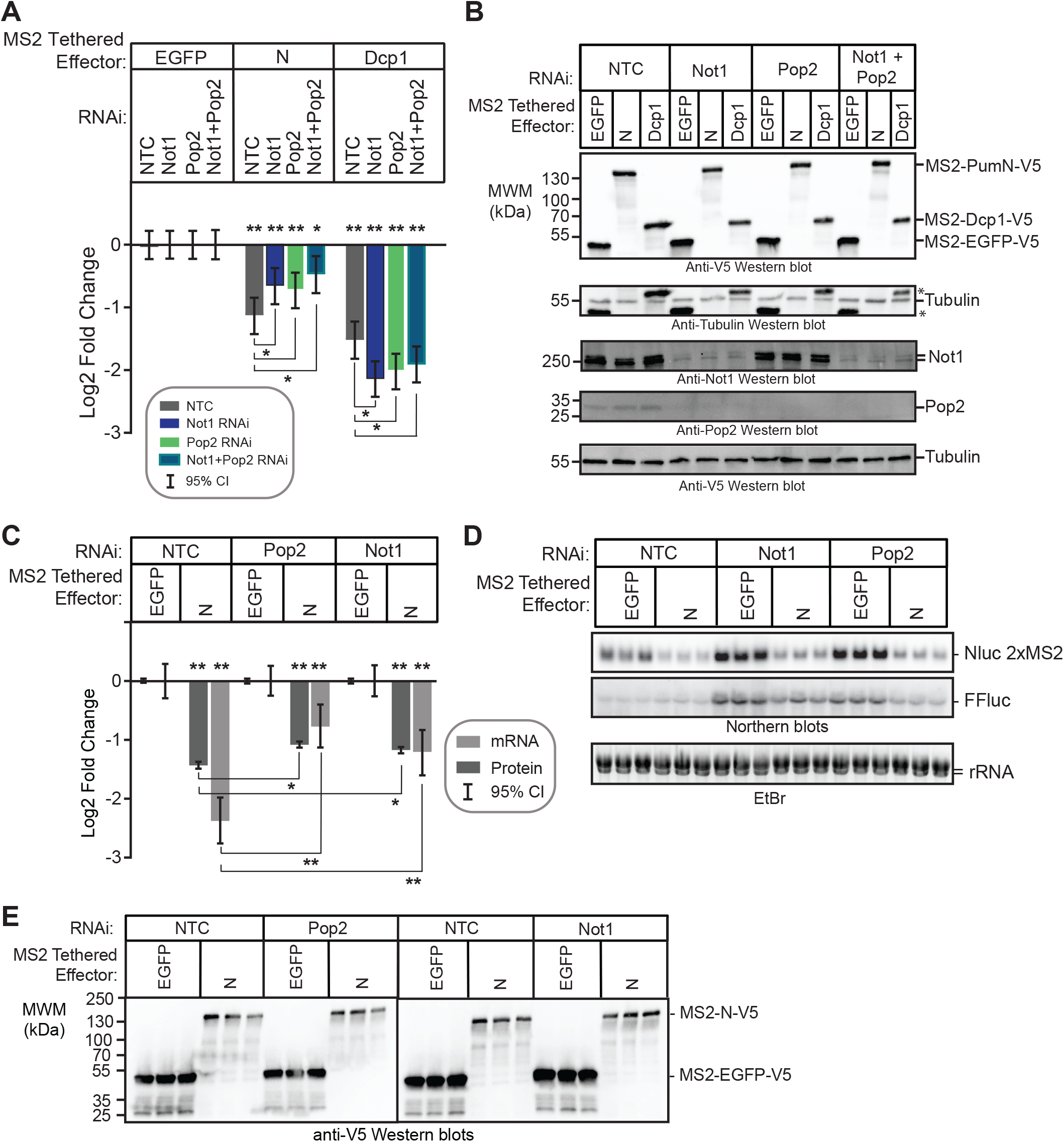
The Pum N-terminus utilizes an additional deadenylation-independent repression activity. (A) The effect of RNAi-mediated depletion of Not1, Pop2 or both simultaneously on the repression activity of the Pum N-terminus was measured in the tethered function reporter assay. The activity of each effector protein was determined relative to the negative control MS2-EGFP within each RNAi condition. Tethered Dcp1 served as a control. The mean log_2_ fold change and 95% credible intervals are graphed for three experimental replicates with four technical replicates each. Data and statistics are reported in Supplementary Table S1. For significance calling, a ‘*’ denotes a posterior probability >0.95 that the difference relative to the negative control is in the indicated direction. The ‘**’ indicates a posterior probability of >0.95 that the indicated difference is at least 1.3-fold. An ‘x’ marks a posterior probability >0.95 that the indicated difference is no more than 1.3-fold in either direction. (B) Western blot of the V5-epitope tagged MS2 fusion effector proteins used in panel A from a representative experimental replicate. Equivalent mass of protein from each sample was probed with anti-V5 antibody, and then with anti-tubulin to assess equal loading of lanes (Asterisks indicate residual anti-V5 signal on the blot). Depletion of Not1 and Pop2 proteins was assessed by Western blot detection of the endogenous proteins. Anti-tubulin Western blot served as a loading control. Note that Not1 depletion also reduces Pop2 protein level, as discussed in the text. (C) The effect of depletion of Pop2 or Not1 on repression activity of Pum N-terminus was measured using the tethered function dual luciferase assay and Northern blotting. Repression of reporter protein and mRNA levels by tethered Pum N-terminus was calculated relative to the negative control MS2-EGFP within the same RNAi condition. Tethered Dcp1 served as a positive control. Mean log_2_ fold change and 95% credible intervals from three biological replicates are reported in the graph. Data and statistics are reported in Supplementary Table S1. (D) Northern blot detection of Nluc 2xMS2 reporter and internal control FFluc mRNAs in the three biological replicates for each effector and RNAi condition. Each lane of the gel contains 5 µg of total RNA. Quantitation of this blot is represented in panel C. Ethidium Bromide detection of rRNA was used to assess integrity and equivalent loading of the RNA samples. (E) Western blot of the V5-tagged MS2 fusion effector proteins used in panel C from three biological replicates. Equivalent mass of protein from each sample was probed with anti-V5 antibody.

**Figure 10.**
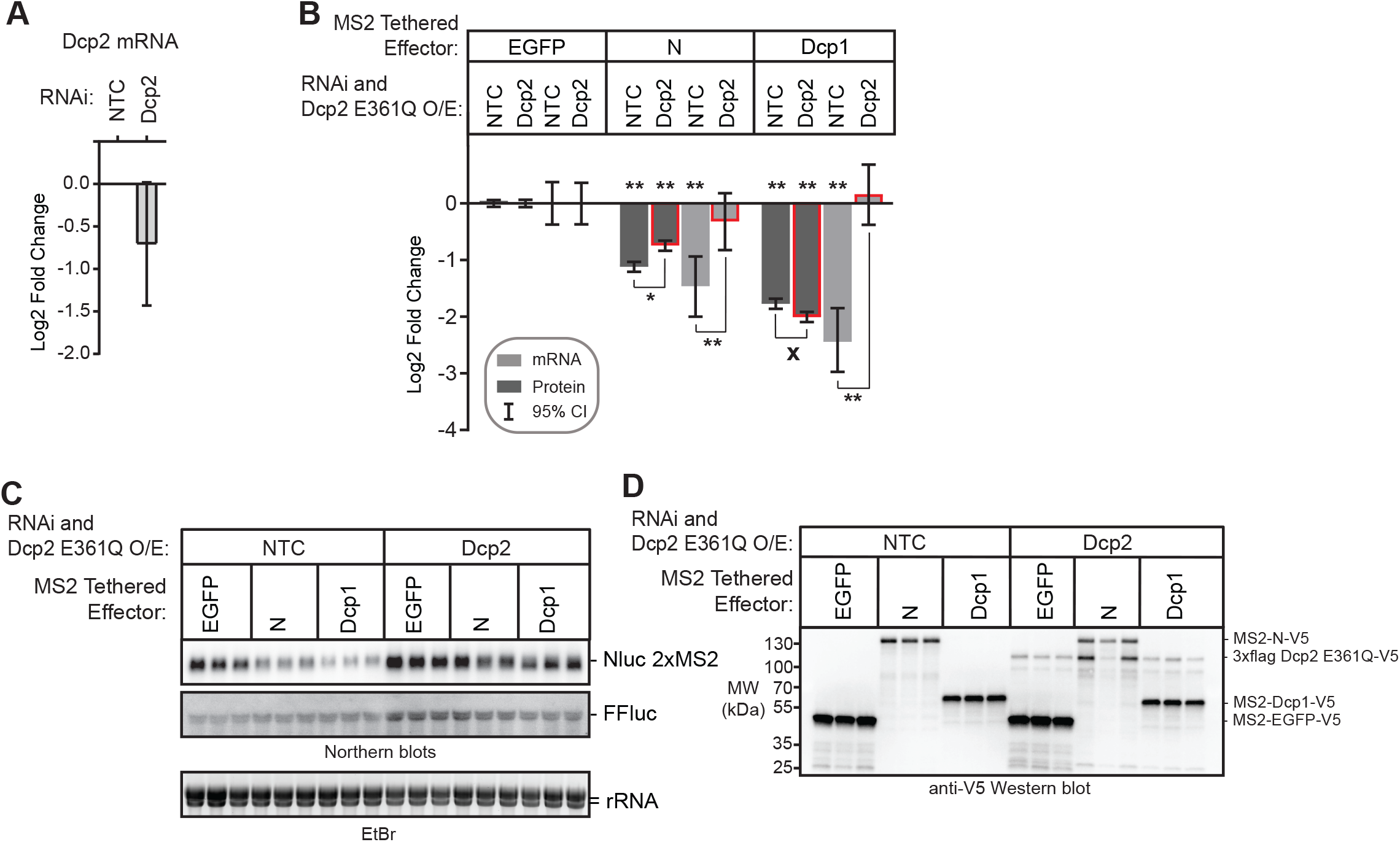
Decapping enzyme participates in repression by the Pum N-terminus. (A) RNAi-mediated depletion of the Dcp2 mRNA after 5 days of dsRNA treatment was measured using RT-qPCR. Fold changes were calculated relative to non-targeting control (NTC). Mean log_2_ fold change and 95% credible intervals for three biological replicates with 3 technical replicates each are reported in the graph. Data and statistics are reported in Supplementary Table S1. (B) The effect of inhibition of decapping on repression of Nluc 2xMS2 reporter protein and mRNA levels by the Pum N-terminus was measured using the tethered function dual luciferase assay. Decapping was inhibited by RNAi mediated depletion of Dcp2 and over-expression of the dominant negative mutant Dcp2 E361Q. Repression activity was calculated relative to tethered EGFP negative control within the same RNAi condition. Tethered Dcp1 served as a positive control. Mean log_2_ fold change and 95% credible intervals are graphed from three biological replicates. Data and statistics are reported in Supplementary Table S1. For significance calling, a ‘*’ denotes a posterior probability >0.95 that the difference relative to the negative control is in the indicated direction. The ‘**’ indicates a posterior probability of >0.95 that the indicated difference is at least 1.3-fold. An ‘x’ marks a posterior probability >0.95 that the indicated difference is no more than 1.3-fold in either direction. (C) Northern blot detection of Nluc 2xMS2 reporter and FFluc internal control in three biological replicate samples for tethered effectors analyzed in panel A. Each lane of the gel contains 5 µg of total RNA. Ethidium Bromide detection of rRNA was used to assess integrity and equivalent loading of the RNA samples. (D) Western blot of the V5-epitope tagged MS2 fusion effector proteins and Dcp2 E361Q used in panel B from three biological replicates. Equivalent mass of protein from each sample was probed with anti-V5 antibody.

How does Pum affect protein synthesis? Because the 5’ cap is crucial for translation of most mRNAs, and the poly(A) tail and pAbp promote translation, Pum-mediated deadenylation and decapping can contribute to both repression of protein synthesis and mRNA destruction. Indeed, Pum-mediated repression of protein level corresponds in magnitude to the reduction in mRNA level. Based on conservation of a cap-binding motif that contains a key tryptophan residue (W783) (48), Pum was proposed to directly inhibit translation by binding the 5’ cap of target mRNAs, and mutation of W783 moderately reduced Pum’s ability to repress a GFP reporter bearing the *mad* 3’UTR in S2 cells (11). However, we did not observe an effect of this mutation on repression by the Pum N-terminus in the tethered function assay or by full length Pum using the PRE reporter (Figure 3). Likewise, our previous analysis showed that the PCMb domain encompassing the putative cap binding motif was neither necessary nor sufficient for repression (36). Thus, the proposed mechanism of cap-dependent inhibition does not appear to make an essential contribution to the Pum activity measured here. We cannot rule out that it might have a potential role in repression in other cellular or developmental contexts, or within the context of certain mRNAs (10, 11).

The Pum RBD contributes to translational repression by associating with and antagonizing the activity of pAbp (44). Consistent with this role, we observed that depletion of pAbp diminished Pum/PRE-mediated repression of protein and mRNA levels (Figure 11). These observations lead us to speculate about the potential functional interplay between Pum, pAbp, and deadenylation. Binding of pAbp to poly(A) was originally thought to interfere with deadenylation (91); however, recent studies indicate that the relationship is more complex (92, 93), wherein Ccr4 and Pop2 deadenylase activities were shown to be differentially affected by pAbp orthologs. Analysis of human PABPC1 and *S. pombe* Pab1 indicate that they are required for Ccr4 deadenylase activity, whereas they inhibit activity of Pop2 orthologs. Further evidence in vitro suggests that *S. pombe* Ccr4 can displace Pab1 from the poly(A) tail, whereas Pop2 cannot. Whether these properties carry over to *Drosophila* remains to be determined. In principle, Pum could cause displacement of pAbp from the mRNA, thereby bypassing the role of Ccr4, resulting in accelerated Pop2-catalyzed deadenylation. Contradicting this hypothesis, however, is the observation based on RNA immunoprecipitation of pAbp, that the Pum RBD did not dislodge pAbp from mRNA (44). Alternatively, the Pum RBD may interfere with the ability of pAbp to promote translation initiation. This idea is supported by functional assays, wherein we showed the Pum RBD inhibits the ability of pAbp to promote translation when bound to an internal poly(A) tract within an mRNA engineered without a 3′ poly(A) tail (44). Future work will include determining the specific interactions between Pum and pAbp, and how this might influence translation efficiency and/or deadenylation.

**Figure 11.**
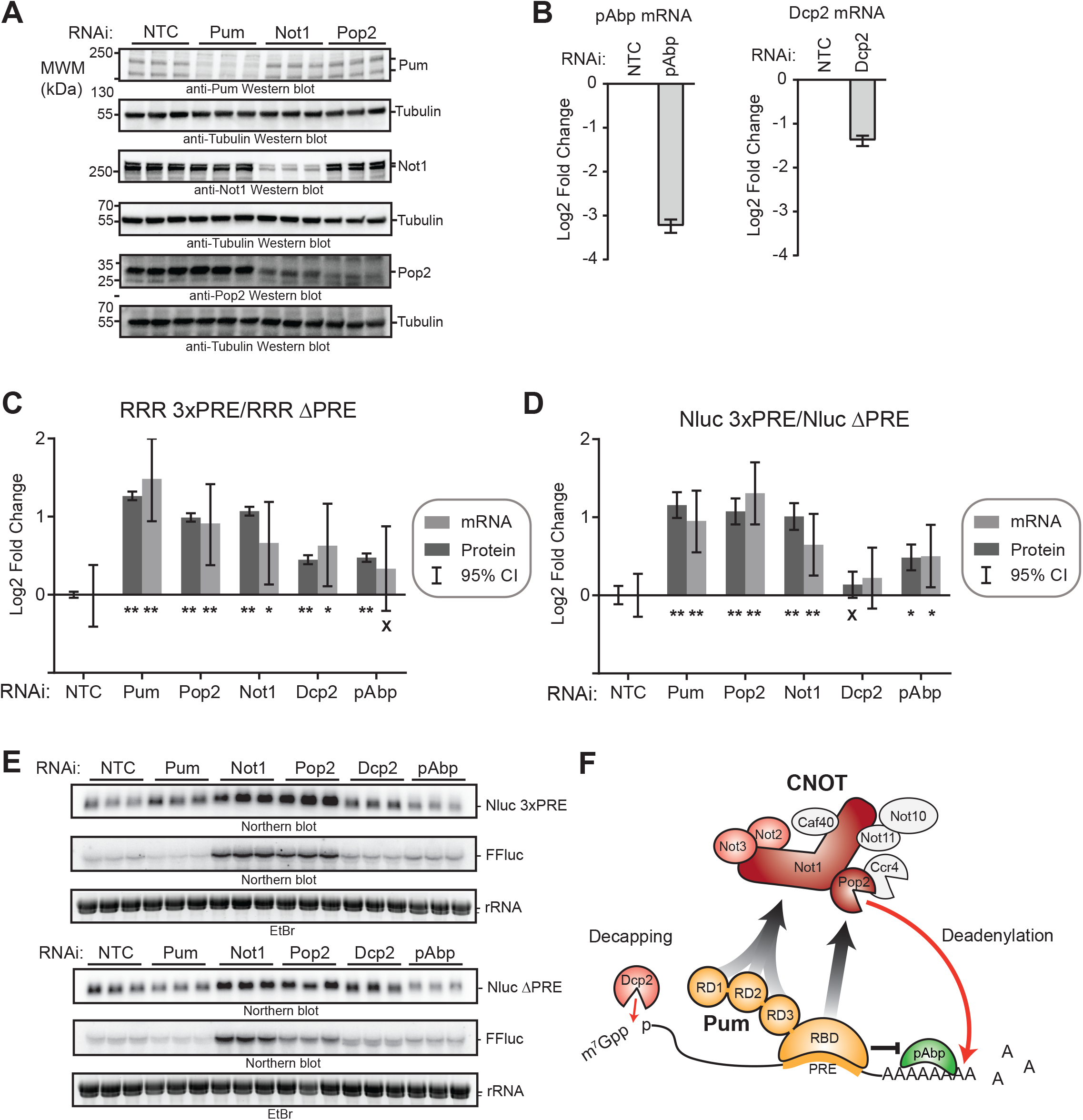
Multiple mechanisms and co-repressors contribute to Pumilio-mediated repression. (A) RNAi-mediated depletion of endogenous Pum, Not1, and Pop2 proteins was assessed by Western blot of three biological replicate samples each from d.mel2 cells that were treated with the indicated dsRNA for three days. Note that Pum and Not1 antibodies each recognize two isoforms of their respective proteins. Equivalent mass of protein was analyzed for each sample, and anti-tubulin Western blots were performed as a loading control. (B) RNAi-mediated depletion of pAbp mRNA (left) and Dcp2 mRNA (right) were measured using RT-qPCR. Fold changes were calculated relative to non-targeting control (NTC). Mean log_2_ fold change and 95% credible intervals for three biological replicates are reported in the graph. Data and statistics are reported in Supplementary Table S1. (C) The effect of RNAi depletion of Pum co-repressors Not1, Pop2, Dcp2, and pAbp on repression of Nluc 3xPRE reporter protein and mRNA expression levels by endogenous Pum was measured in d.mel-2 cells. Data was analyzed by calculating the Relative Response Ratio for each sample by dividing the Nluc signal by corresponding FFluc signal, thereby normalizing variation in transfection efficiency. Next, the PRE-dependent effect of each RNAi condition on the Pum repressed, PRE containing reporter was normalized to the effect on the unregulated Nluc ΔPRE reporter, which contains a minimal 3’UTR that lacks Pum binding sites. The fold change in PRE-mediated regulation within each RNAi condition was then calculated relative to the negative control NTC dsRNA. RNAi of Pum served as a positive control. Mean log_2_ fold change and 95% credible intervals for three biological replicates are reported in the graphs. Data and statistics are reported in Supplementary Table S1. For significance calling, a ‘*’ denotes a posterior probability >0.95 that the difference relative to the negative control is in the indicated direction. The ‘**’ indicates a posterior probability of >0.95 that the indicated difference is at least 1.3-fold. An ‘x’ marks a posterior probability >0.95 that the indicated difference is no more than 1.3-fold in either direction. (D) The effect of RNAi depletion of Pum co-repressors Not1, Pop2, Dcp2, and pAbp on repression of Nluc 3xPRE reporter protein and mRNA expression levels by endogenous Pum was analyzed as in Panel C except that the FFluc protein and mRNA values were omitted. The PRE-dependent effect of each RNAi condition on the Nluc 3xPRE reporter was normalized to the effect on the unregulated Nluc ΔPRE reporter. The fold change in PRE-mediated regulation within each RNAi condition was then calculated relative to the negative control NTC dsRNA. RNAi of Pum served as a positive control. Mean log_2_ fold change and 95% credible intervals for three biological replicates are reported in the graphs. Data and statistics are reported in Supplementary Table S1. (E) Northern blot detection of Pum-regulated Nluc 3xPRE, unregulated Nluc ΔPRE reporter mRNA and FFluc internal control mRNAs in three biological replicate samples for each RNAi condition analyzed in panel C and D. Each lane of the gel contains 5 µg of total RNA. Ethidium Bromide detection of rRNA was used to assess integrity and equivalent loading of the RNA samples. (F) Model of Pum-mediated repression. The RNA-binding domain (RBD) of Pum binds to mRNAs that contain a Pum Response Element (PRE). Multiple domains of Pum contribute to repression activity including the N-terminal repression domains (RD1, RD2, and RD3) and C-terminal RBD. Pum represses the target mRNA by multiple mechanisms including acceleration of mRNA decay via recruitment of Ccr4-Pop2-Not (CNOT) deadenylase complex, leading to deadenylation of the 3’ poly-adenosine tail, and via decapping enzyme (Dcp2) mediated removal of the 5’ 7-methyl guanosine cap (7mGppp). Pum RBD also antagonizes the translational activity of poly-adenosine binding protein (pAbp). CNOT subunits that are important for Pum RD-mediated repression are shaded in red. Red arrows indicate enzyme-catalyzed hydrolysis of the RNA. Arrows with grey-black gradient indicate Pum-CNOT interactions, as described in the Discussion. The means by which Dcp2 is modulated by Pum N-terminus remains to be determined.

Our results have important implications for understanding Pum-mediated repression in embryos, the germline, and neurons. Pum repression of *hunchback* mRNA in early embryogenesis was linked to poly(A) tail shortening (8). Pum repression was also correlated with mRNA decay during the maternal-to-zygotic transition (MZT), wherein many PRE-containing, maternally provided mRNAs are coordinately degraded (28,29,94). Pum-mediated repression was also linked to deadenylation by CNOT in primordial germ cells, where Pum contributes to repression of *cyclin B* mRNA (42), and in GSCs, where Pum participates in repression of mRNAs such as *mei-P26* (43). The mechanisms of Pum-mediated deadenylation and decapping likely contribute to mRNA degradation observed in these contexts. Pum also represses specific mRNAs in neurons, such as *paralytic*, which encodes a voltage-gated sodium channel (15,16,95); however, the impact on mRNA decay remains to be examined in this context. Future analysis should investigate the contributions of Pum RDs to these and other processes.

Regulation of a growing number of Pum target mRNAs involves collaboration with other RBPs, with Nanos and Brat being the best documented examples. How can the mechanisms of Pum repression integrate with the activities of these RBPs? In the case of Nanos, it binds cooperatively with Pum to certain target mRNAs that possess a Nanos binding site preceding a PRE motif (25, 96). Nanos also confers its own repressive activity that accelerates deadenylation (45). Nanos may synergize with Pum in the recruitment of CNOT by contributing additional contacts with the Not1 and Not3 subunits. Nanos was also reported to interact with Not4 (42), though the potential role of Not4 in deadenylation and its involvement in Nanos activity are unclear. In fact, Drosophila Not4 does not appear to be a stable, stoichiometric CNOT component (40). Nanos also promotes decapping, but like Pum, how it does so is presently not well understood (45). Thus, combinatorial control by Pum and Nanos together accelerate the same key steps contributing to silencing of translation and mRNA destruction (2).

Pum can also collaborate with Brat to repress certain mRNAs that contain both a PRE and Brat binding site (10,11,28,33,34). The mechanism of Brat-mediated repression in embryos was reported to involve recruitment of the translational repressor eIF4E homologous protein (4EHP) (32). Brat may also affect mRNA decay, supported by the observation that it co-purifies with the CNOT complex (40). Depletion of Pop2 reduced the combined repressive activity of Pum and Brat (11). Future analyses will be necessary to interrogate the role of CNOT in Brat-mediated repression and whether Pum and Brat can synergistically recruit CNOT to their mutual target mRNAs. Pum, Nanos, and Brat all collaborate to bind and repress the maternal *hunchback* mRNA during embryogenesis through a combination of cooperative RNA-binding, translational repression, deadenylation, and mRNA decay (reviewed in (2)). It is noteworthy that the assembly of this triumvirate of repressors on one mRNA is likely a rare scenario, as few mRNAs are predicted to contain the requisite cluster of binding sites for these three RBPs (2).

The mechanistic insights into Pum-mediated repression also have important implications for Pum orthologs in other species. *Drosophila* Pum serves as an archetype, and the conservation of its repressive domains, including those in its N-terminus and RBD, in animals ranging from insects to vertebrates indicates that the mechanisms and co-repressors described here will be relevant (47). Indeed, as noted above, accumulating evidence supports the role of deadenylation, decapping, and translational inhibition in Pum repression in multiple model organisms. Mammalian Pum orthologs have crucial, diverse roles in growth and development, gametogenesis, hematopoiesis, neurogenesis, behavior, motor function and memory formation (97–106). Their dysfunction has now been linked to cancer, neurodegeneration, epilepsy, memory impairment, reduced fertility, and developmental defects in mammals (97,98,100,103,107–111). Thus, we anticipate that further elucidation of Pum function will facilitate better understanding of its role in development and disease, and perhaps inform future therapeutic interventions.

## ACKNOWLEDGMENTS

We thank Dr. Elmar Wahle for providing CNOT antibodies and Dr. Eric Wagner for sharing the pUB plasmid and IntS1 antibody. We also thank the members of the Goldstrohm lab for helpful advice, including Katherine McKenney, Isioma Enwerem, and Elizabeth Abshire for editing assistance.

## COMPETING INTERESTS

None

**Supplementary Table 1**

Tables of all data, number and type of replicates, and statistical values are reported for all figures. Data is organized by tabs corresponding to each figure.

**Supplementary File 1**

Tables of all oligonucleotides, plasmids, reagents, and antibodies used in this study are provided, along with relevant experimental parameters and features.

**Supplementary File 2**

Checklist of minimum information for publication of quantitative real-time PCR experiments (MIQE), along with details of qPCR primers, their target genes, amplicons, and optimization.

---

**Supplementary Figure S1.**
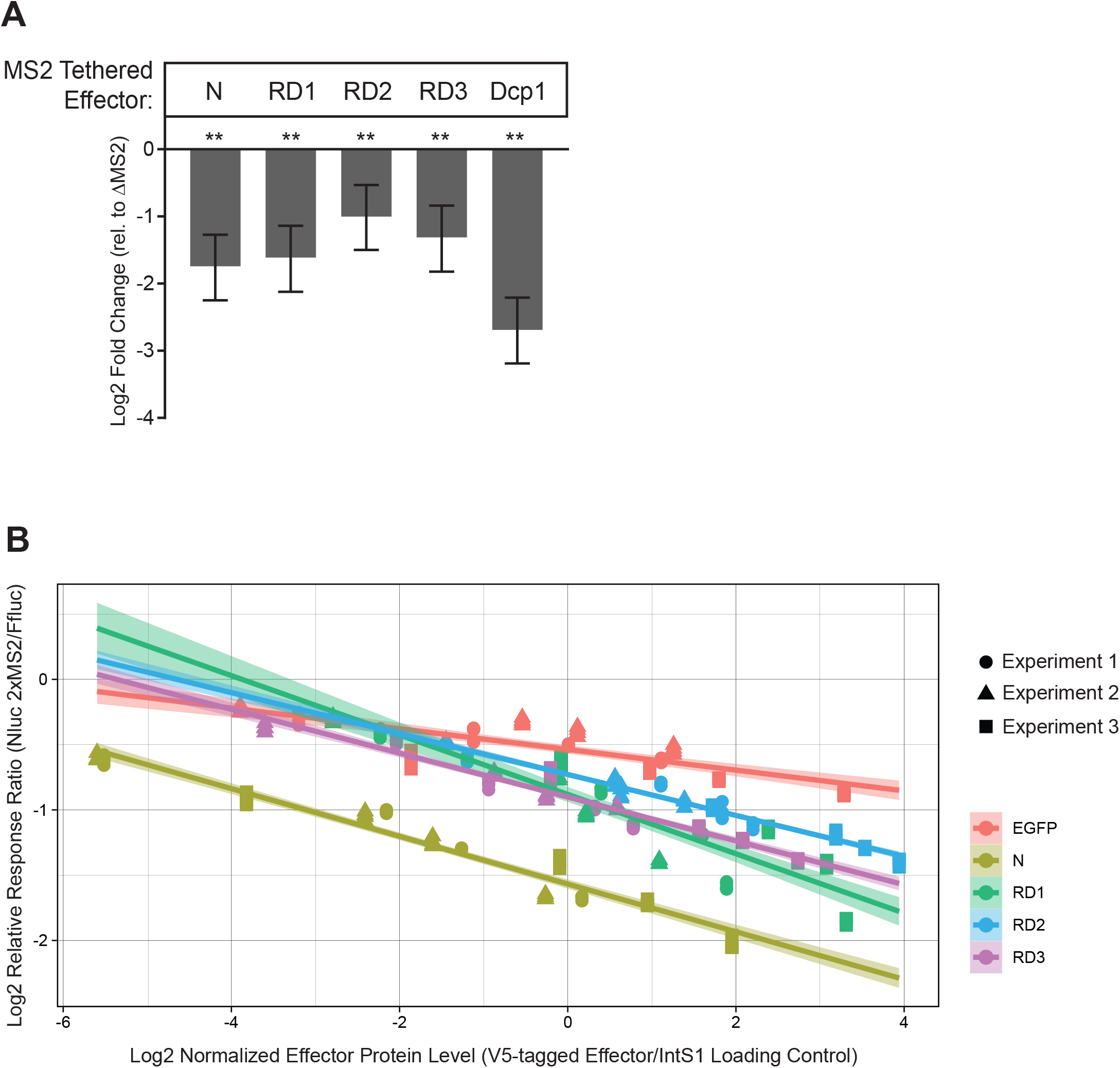
Repression by tethered effectors exhibits a log-linear relationship to effector protein level. (A) The specificity of the repression activity of tethered effector proteins (Pum N-terminus, individual repression domains RD1, RD2, RD3, and Dcp1) were measured using the tethered function dual luciferase assay using the Nluc 2xMS2 reporter or an equivalent reporter wherein the MS2 binding sites are deleted, Nluc ΔMS2. The data are the same as in Figure 2B, except that the mean log_2_ fold change of normalized Nluc 2xMS2 reporter activity was calculated relative to Nluc ΔMS2. Mean fold change values are plotted with 95% credible intervals from 3 experiments with 4 technical replicates each. Data and statistics are reported in Supplementary Table S1. For significance calling, the ‘**’ indicates a posterior probability of >0.95 that the indicated difference is at least 1.3-fold. (B) The relationship of repression activity to the level of expressed protein for titrations of each MS2 tethered effector was measured using the tethered function assay (data from Figure 2D). The amount of expressed V5-tagged effector proteins were measured by quantitative western blotting (see Figure 2E for representative Western blot) and normalized to the amount of the IntS1 loading control for each sample. The reporter activity of Nluc 2xMS2 was divided to internal control FFluc signal for each sample to calculate the Relative Response Ratio, so as to normalize variations in transfection efficiency. The log_2_ Relative Response Ratios from three experiments, each with four technical replicates, are plotted versus the log_2_ normalized effector protein level from the three experiments. The associated linear trend lines (bold lines) show the log-linear relationship of the activities of each effector per unit protein. The shapes (circle, triangle, and square) represent data points from each of the experimental replicates. Shaded regions show the 95% confidence intervals for the trend lines. Data and statistics are reported in Supplementary Table S1.

**Supplementary Figure S2.**
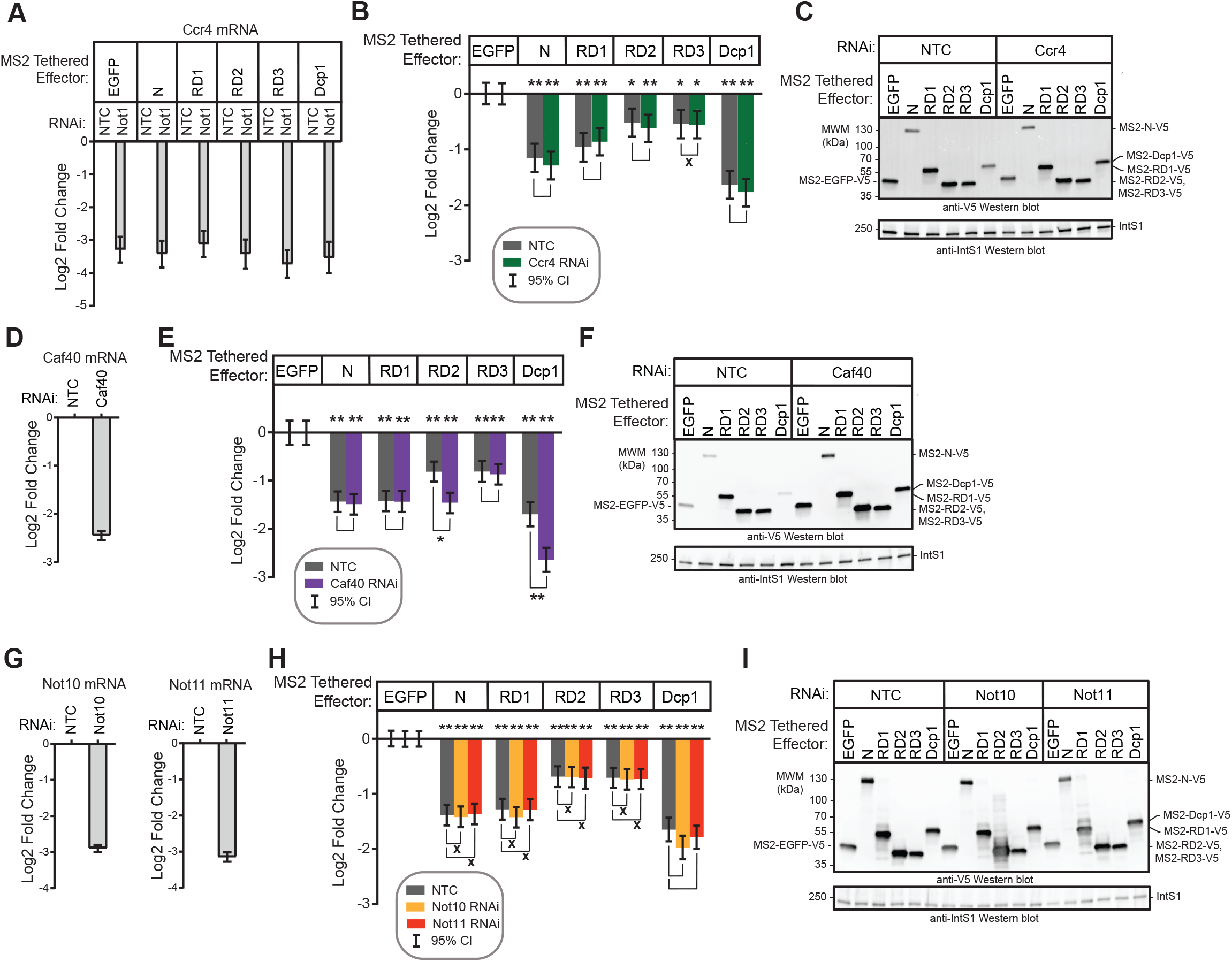
Depletion of Ccr4, Caf40, Not10 and Not11 does not impair tethered Pum repression activity. (A) Depletion of Ccr4 mRNA by RNAi after 3 days of treatment with the indicated dsRNA was measured by RT-qPCR. Fold changes were calculated relative to non-targeting control (NTC). Mean log_2_ fold change and 95% credible intervals are reported in the graph for three biological replicates with three technical replicates each. Data and statistics are reported in Supplementary Table S1. (B) Tethered function dual luciferase assays measured the effect of Ccr4 depletion on the repression activity of Pum N-terminus and RDs. Non-targeting control (NTC) serves as negative control for comparison. Activity of each effector was determined relative to tethered EGFP negative control in each RNAi condition. Mean log_2_ fold change and 95% credible intervals are reported in the graph from three experimental replicates with four technical replicates each. Data and statistics are reported in Supplementary Table S1. For significance calling, a ‘*’ denotes a posterior probability >0.95 that the difference relative to the negative control is in the indicated direction. The ‘**’ indicates a posterior probability of >0.95 that the indicated difference is at least 1.3-fold. An ‘x’ marks a posterior probability >0.95 that the indicated difference is no more than 1.3-fold in either direction. (C) Western blot of the V5-epitope tagged MS2 fusion effector proteins used in panel B from a representative experimental replicate. Equivalent mass of protein from each sample was probed with anti-V5 antibody, followed by anti-IntS1 western blot as a loading control. (D) RT-qPCR was used to measure the efficiency of Caf40 mRNA depletion after 3 days of dsRNA treatment. Fold changes were calculated relative to non-targeting control (NTC). Mean log_2_ fold change and 95% credible intervals are reported in the graph for three biological replicates with three technical replicates each. Data and statistics are reported in Supplementary Table S1. (E) Tethered function dual luciferase assays measured the effect of Caf40 depletion on the repression activity of Pum N-terminus and RDs. Non-targeting control (NTC) serves as negative control for comparison. Activity of each effector was determined relative to tethered EGFP negative control in each RNAi condition. Mean log_2_ fold change and 95% credible intervals are reported in the graph for three experimental replicates with four technical replicates each. Data and statistics are reported in Supplementary Table S1. (F) Western blot of the V5-epitope tagged MS2 fusion effector proteins used in panel E from a representative experimental replicate. Equivalent mass of protein from each sample was probed with anti-V5 antibody, followed by anti-IntS1 western blot as a loading control. (G) RT-qPCR was used to measure the efficiency of Not10 (left) and Not11 (right) mRNA depletion after 3 days of dsRNA treatment. Fold changes were calculated relative to non-targeting control (NTC). Mean log_2_ fold change and 95% credible intervals are reported in the graph for three biological replicates with three technical replicates each. Data and statistics are reported in Supplementary Table S1. (H) Tethered function dual luciferase assays measured the effect of Not10 and Not11 depletion on the repression activity of Pum N-terminus and RDs. Non-targeting control (NTC) serves as negative control for comparison. Activity of each effector was determined relative to tethered EGFP negative control in each RNAi condition. Mean log_2_ fold change and 95% credible intervals are reported in the graph for three experimental replicates with four technical replicates each. Data and statistics are reported in Supplementary Table S1. (I) Western blot of the V5-epitope tagged MS2 fusion effector proteins used in panel H from a representative experimental replicate. Equivalent mass of protein from each sample was probed with anti-V5 antibody, followed by anti-IntS1 western blot as a loading control.

**Supplementary Figure S3.**
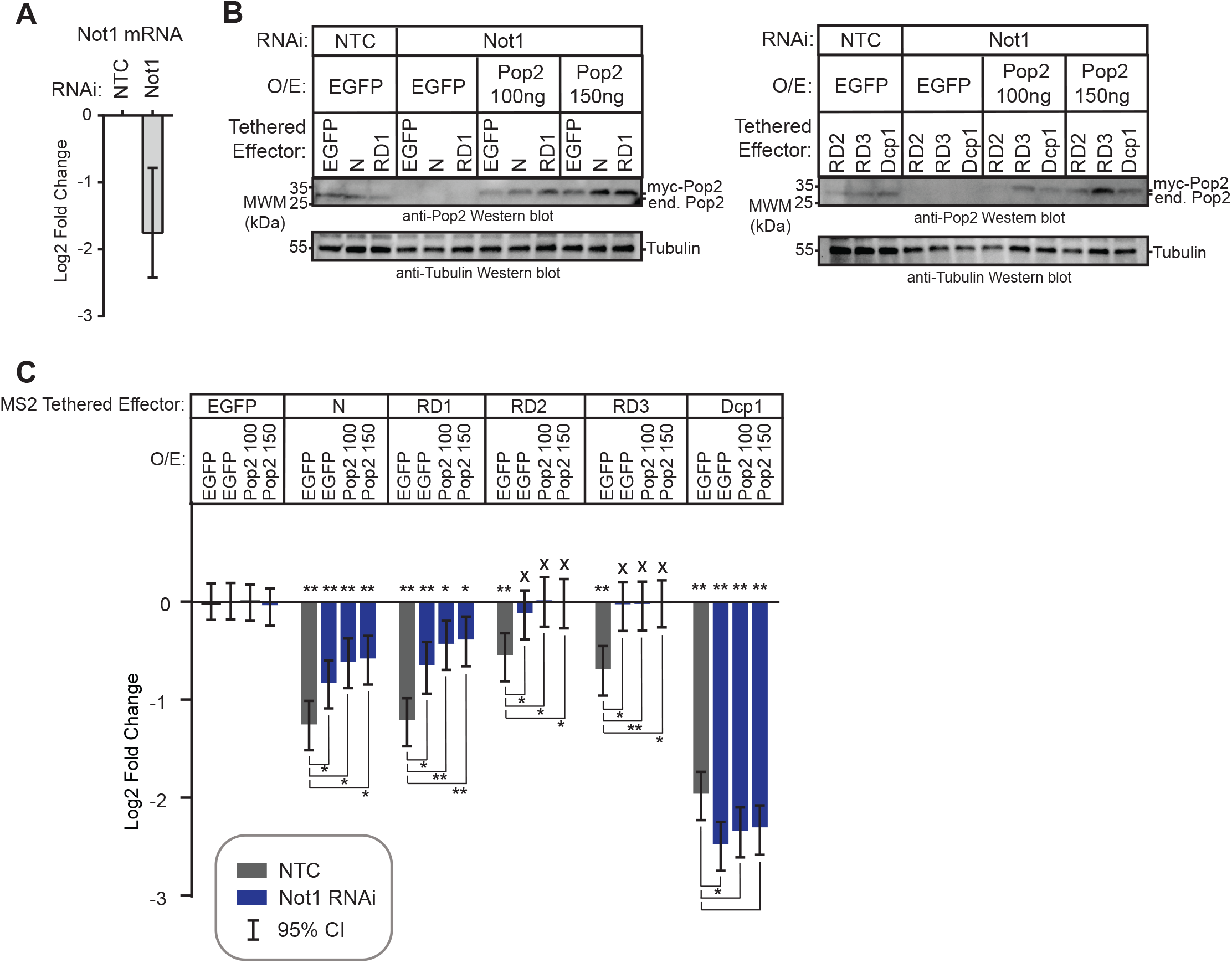
Pop2 expression does not rescue impaired Pum N-terminus and RD activity from Not1 depletion. (A) The efficiency of Not1 mRNA depletion after 5 days of dsRNA treatment was measured using RT-qPCR. Fold changes were calculated relative to non-targeting control (NTC). Mean log_2_ fold change and 95% credible intervals from three biological replicates are reported in the graph. Data and statistics are reported in Supplementary Table S1. (B) Western blots confirming depletion of endogenous Pop2 protein resulting from RNAi-mediated depletion of Not1. Rescue of Pop2 included two amounts of transfected myc-Pop2 plasmid (100ng or 150ng) to restore Pop2 to approximately endogenous levels. Equivalent amount of protein in each cell extract was analyzed by western blot. Tubulin western blot serves as a loading control. (C) The ability of Pop2 to rescue Pum RD mediated repression in Not1 depleted cells was analyzed using the tethered function dual luciferase assay. The effect of RNAi-mediated depletion of Not1 on repression by Pum N-terminus and RDs relative to tethered EGFP was measured. Myc-Pop2 was expressed using 100ng and 150ng of transfected plasmid. Mean log_2_ fold change and 95% credible intervals are reported in the graph for four experimental replicates with four technical replicates each. For significance calling, a ‘*’ denotes a posterior probability >0.95 that the difference relative to the negative control is in the indicated direction. The ‘**’ indicates a posterior probability of >0.95 that the indicated difference is at least 1.3-fold. An ‘x’ marks a posterior probability >0.95 that the indicated difference is no more than 1.3-fold in either direction.

## REFERENCES

1. Wickens, M., Bernstein, D.S., Kimble, J. and Parker, R. (2002) A PUF family portrait: 3’UTR regulation as a way of life. Trends Genet, 18, 150–157.

2. Arvola, R.M., Weidmann, C.A., Tanaka Hall, T.M. and Goldstrohm, A.C. (2017) Combinatorial control of messenger RNAs by Pumilio, Nanos and Brain Tumor Proteins. RNA Biol, 14, 1445–1456.

3. Lehmann, R. and Nusslein-Volhard, C. (1987) Involvement of the pumilio gene in the transport of an abdominal signal in the Drosophila embyro. Nature, 329, 167–170.

4. Barker, D.D., Wang, C., Moore, J., Dickinson, L.K. and Lehmann, R. (1992) Pumilio is essential for function but not for distribution of the Drosophila abdominal determinant Nanos. Genes Dev, 6, 2312–2326.

5. Murata, Y. and Wharton, R.P. (1995) Binding of pumilio to maternal hunchback mRNA is required for posterior patterning in Drosophila embryos. Cell, 80, 747–756.

6. Struhl, G., Johnston, P. and Lawrence, P.A. (1992) Control of Drosophila body pattern by the hunchback morphogen gradient. Cell, 69, 237–249.

7. Lehmann, R. and Nusslein-Volhard, C. (1987) hunchback, a gene required for segmentation of an anterior and posterior region of the Drosophila embryo. Dev Biol, 119, 402–417.

8. Wreden, C., Verrotti, A.C., Schisa, J.A., Lieberfarb, M.E. and Strickland, S. (1997) Nanos and pumilio establish embryonic polarity in Drosophila by promoting posterior deadenylation of hunchback mRNA. Development, 124, 3015–3023.

9. Irish, V., Lehmann, R. and Akam, M. (1989) The Drosophila posterior-group gene nanos functions by repressing hunchback activity. Nature, 338, 646–648.

10. Harris, R.E., Pargett, M., Sutcliffe, C., Umulis, D. and Ashe, H.L. (2011) Brat promotes stem cell differentiation via control of a bistable switch that restricts BMP signaling. Dev Cell, 20, 72–83.

11. Newton, F.G., Harris, R.E., Sutcliffe, C. and Ashe, H.L. (2015) Coordinate post-transcriptional repression of Dpp-dependent transcription factors attenuates signal range during development. Development, 142, 3362–3373.

12. Forbes, A. and Lehmann, R. (1998) Nanos and Pumilio have critical roles in the development and function of Drosophila germline stem cells. Development, 125, 679–690.

13. Lin, H. and Spradling, A.C. (1997) A novel group of pumilio mutations affects the asymmetric division of germline stem cells in the Drosophila ovary. Development, 124, 2463–2476.

14. Asaoka-Taguchi, M., Yamada, M., Nakamura, A., Hanyu, K. and Kobayashi, S. (1999) Maternal Pumilio acts together with Nanos in germline development in Drosophila embryos. Nat Cell Biol, 1, 431–437.

15. Mee, C.J., Pym, E.C., Moffat, K.G. and Baines, R.A. (2004) Regulation of neuronal excitability through pumilio-dependent control of a sodium channel gene. J Neurosci, 24, 8695–8703.

16. Muraro, N.I., Weston, A.J., Gerber, A.P., Luschnig, S., Moffat, K.G. and Baines, R.A. (2008) Pumilio binds para mRNA and requires Nanos and Brat to regulate sodium current in Drosophila motoneurons. J Neurosci, 28, 2099–2109.

17. Bhogal, B., Plaza-Jennings, A. and Gavis, E.R. (2016) Nanos-mediated repression of hid protects larval sensory neurons after a global switch in sensitivity to apoptotic signals. Development, 143, 2147–2159.

18. Ye, B., Petritsch, C., Clark, I.E., Gavis, E.R., Jan, L.Y. and Jan, Y.N. (2004) Nanos and Pumilio are essential for dendrite morphogenesis in Drosophila peripheral neurons. Curr Biol, 14, 314–321.

19. Dubnau, J., Chiang, A.S., Grady, L., Barditch, J., Gossweiler, S., McNeil, J., Smith, P., Buldoc, F., Scott, R., Certa, U. et al. (2003) The staufen/pumilio pathway is involved in Drosophila long-term memory. Curr Biol, 13, 286–296.

20. Menon, K.P., Sanyal, S., Habara, Y., Sanchez, R., Wharton, R.P., Ramaswami, M. and Zinn, K. (2004) The translational repressor Pumilio regulates presynaptic morphology and controls postsynaptic accumulation of translation factor eIF-4E. Neuron, 44, 663–676.

21. Menon, K.P., Andrews, S., Murthy, M., Gavis, E.R. and Zinn, K. (2009) The translational repressors Nanos and Pumilio have divergent effects on presynaptic terminal growth and postsynaptic glutamate receptor subunit composition. J Neurosci, 29, 5558–5572.

22. Zamore, P.D., Bartel, D.P., Lehmann, R. and Williamson, J.R. (1999) The PUMILIO-RNA interaction: a single RNA-binding domain monomer recognizes a bipartite target sequence. Biochemistry, 38, 596–604.

23. Zamore, P.D., Williamson, J.R. and Lehmann, R. (1997) The Pumilio protein binds RNA through a conserved domain that defines a new class of RNA-binding proteins. Rna, 3, 1421–1433.

24. Lou, T.F., Weidmann, C.A., Killingsworth, J., Tanaka Hall, T.M., Goldstrohm, A.C. and Campbell, Z.T. (2016) Integrated analysis of RNA-binding protein complexes using in vitro selection and high-throughput sequencing and sequence specificity landscapes (SEQRS). Methods.

25. Weidmann, C.A., Qiu, C., Arvola, R.M., Lou, T.F., Killingsworth, J., Campbell, Z.T., Tanaka Hall, T.M. and Goldstrohm, A.C. (2016) Drosophila Nanos acts as a molecular clamp that modulates the RNA-binding and repression activities of Pumilio. Elife, 5.

26. Wang, X., McLachlan, J., Zamore, P.D. and Hall, T.M. (2002) Modular recognition of RNA by a human pumilio-homology domain. Cell, 110, 501–512.

27. Gerber, A.P., Luschnig, S., Krasnow, M.A., Brown, P.O. and Herschlag, D. (2006) Genome-wide identification of mRNAs associated with the translational regulator PUMILIO in Drosophila melanogaster. Proc Natl Acad Sci U S A, 103, 4487–4492.

28. Laver, J.D., Li, X., Ray, D., Cook, K.B., Hahn, N.A., Nabeel-Shah, S., Kekis, M., Luo, H., Marsolais, A.J., Fung, K.Y. et al. (2015) Brain tumor is a sequence-specific RNA-binding protein that directs maternal mRNA clearance during the Drosophila maternal-to-zygotic transition. Genome Biol, 16, 94.

29. Laver, J.D., Marsolais, A.J., Smibert, C.A. and Lipshitz, H.D. (2015) Regulation and Function of Maternal Gene Products During the Maternal-to-Zygotic Transition in Drosophila. Curr Top Dev Biol, 113, 43–84.

30. Sonoda, J. and Wharton, R.P. (2001) Drosophila Brain Tumor is a translational repressor. Genes Dev, 15, 762–773.

31. Edwards, T.A., Wilkinson, B.D., Wharton, R.P. and Aggarwal, A.K. (2003) Model of the brain tumor-Pumilio translation repressor complex. Genes Dev, 17, 2508–2513.

32. Cho, P.F., Gamberi, C., Cho-Park, Y.A., Cho-Park, I.B., Lasko, P. and Sonenberg, N. (2006) Cap-dependent translational inhibition establishes two opposing morphogen gradients in Drosophila embryos. Curr Biol, 16, 2035–2041.

33. Loedige, I., Jakob, L., Treiber, T., Ray, D., Stotz, M., Treiber, N., Hennig, J., Cook, K.B., Morris, Q., Hughes, T.R. et al. (2015) The Crystal Structure of the NHL Domain in Complex with RNA Reveals the Molecular Basis of Drosophila Brain-Tumor-Mediated Gene Regulation. Cell Rep, 13, 1206–1220.

34. Loedige, I., Stotz, M., Qamar, S., Kramer, K., Hennig, J., Schubert, T., Loffler, P., Langst, G., Merkl, R., Urlaub, H. et al. (2014) The NHL domain of BRAT is an RNA-binding domain that directly contacts the hunchback mRNA for regulation. Genes Dev, 28, 749–764.

35. Wharton, R.P. and Struhl, G. (1991) RNA regulatory elements mediate control of Drosophila body pattern by the posterior morphogen nanos. Cell, 67, 955–967.

36. Weidmann, C.A. and Goldstrohm, A.C. (2012) Drosophila Pumilio protein contains multiple autonomous repression domains that regulate mRNAs independently of Nanos and brain tumor. Mol Cell Biol, 32, 527–540.

37. Jackson, R.J., Hellen, C.U. and Pestova, T.V. (2010) The mechanism of eukaryotic translation initiation and principles of its regulation. Nat Rev Mol Cell Biol, 11, 113–127.

38. Garneau, N.L., Wilusz, J. and Wilusz, C.J. (2007) The highways and byways of mRNA decay. Nat Rev Mol Cell Biol, 8, 113–126.

39. Goldstrohm, A.C. and Wickens, M. (2008) Multifunctional deadenylase complexes diversify mRNA control. Nat Rev Mol Cell Biol, 9, 337–344.

40. Temme, C., Zhang, L., Kremmer, E., Ihling, C., Chartier, A., Sinz, A., Simonelig, M. and Wahle, E. (2010) Subunits of the Drosophila CCR4-NOT complex and their roles in mRNA deadenylation. Rna, 16, 1356–1370.

41. Temme, C., Zaessinger, S., Meyer, S., Simonelig, M. and Wahle, E. (2004) A complex containing the CCR4 and CAF1 proteins is involved in mRNA deadenylation in Drosophila. Embo J, 23, 2862–2871.

42. Kadyrova, L.Y., Habara, Y., Lee, T.H. and Wharton, R.P. (2007) Translational control of maternal Cyclin B mRNA by Nanos in the Drosophila germline. Development, 134, 1519–1527.

43. Joly, W., Chartier, A., Rojas-Rios, P., Busseau, I. and Simonelig, M. (2013) The CCR4 deadenylase acts with Nanos and Pumilio in the fine-tuning of Mei-P26 expression to promote germline stem cell self-renewal. Stem Cell Reports, 1, 411–424.

44. Weidmann, C.A., Raynard, N.A., Blewett, N.H., Van Etten, J. and Goldstrohm, A.C. (2014) The RNA binding domain of Pumilio antagonizes poly-adenosine binding protein and accelerates deadenylation. RNA, 20, 1298–1319.

45. Raisch, T., Bhandari, D., Sabath, K., Helms, S., Valkov, E., Weichenrieder, O. and Izaurralde, E. (2016) Distinct modes of recruitment of the CCR4-NOT complex by Drosophila and vertebrate Nanos. EMBO J, 35, 974–990.

46. Bhandari, D., Raisch, T., Weichenrieder, O., Jonas, S. and Izaurralde, E. (2014) Structural basis for the Nanos-mediated recruitment of the CCR4-NOT complex and translational repression. Genes Dev, 28, 888–901.

47. Goldstrohm, A.C., Hall, T.M.T. and McKenney, K.M. (2018) Post-transcriptional Regulatory Functions of Mammalian Pumilio Proteins. Trends Genet, 34, 972–990.

48. Cao, Q., Padmanabhan, K. and Richter, J.D. (2010) Pumilio 2 controls translation by competing with eIF4E for 7-methyl guanosine cap recognition. Rna, 16, 221–227.

49. Wharton, R.P., Sonoda, J., Lee, T., Patterson, M. and Murata, Y. (1998) The Pumilio RNA-binding domain is also a translational regulator. Mol Cell, 1, 863–872.

50. Oh, S.K., Scott, M.P. and Sarnow, P. (1992) Homeotic gene Antennapedia mRNA contains 5’-noncoding sequences that confer translational initiation by internal ribosome binding. Genes Dev, 6, 1643–1653.

51. Ye, X., Fong, P., Iizuka, N., Choate, D. and Cavener, D.R. (1997) Ultrabithorax and Antennapedia 5’ untranslated regions promote developmentally regulated internal translation initiation. Mol Cell Biol, 17, 1714–1721.

52. Nishihara, T., Zekri, L., Braun, J.E. and Izaurralde, E. (2013) miRISC recruits decapping factors to miRNA targets to enhance their degradation. Nucleic Acids Res, 41, 8692–8705.

53. Haas, G., Braun, J.E., Igreja, C., Tritschler, F., Nishihara, T. and Izaurralde, E. (2010) HPat provides a link between deadenylation and decapping in metazoa. J Cell Biol, 189, 289–302.

54. Gibson, D.G., Young, L., Chuang, R.Y., Venter, J.C., Hutchison, C.A., 3rd and Smith, H.O. (2009) Enzymatic assembly of DNA molecules up to several hundred kilobases. Nat Methods, 6, 343-345.

55. Diebold, M.L., Fribourg, S., Koch, M., Metzger, T. and Romier, C. (2011) Deciphering correct strategies for multiprotein complex assembly by co-expression: application to complexes as large as the histone octamer. J Struct Biol, 175, 178–188.

56. Harlow, E. and Lane, D. (2006) Purification of antibodies on an antigen column. CSH Protoc, 2006.

57. Raisch, T., Chang, C.T., Levdansky, Y., Muthukumar, S., Raunser, S. and Valkov, E. (2019) Reconstitution of recombinant human CCR4-NOT reveals molecular insights into regulated deadenylation. Nat Commun, 10, 3173.

58. Lai, W.S., Arvola, R.M., Goldstrohm, A.C. and Blackshear, P.J. (2019) Inhibiting transcription in cultured metazoan cells with actinomycin D to monitor mRNA turnover. Methods, 155, 77–87.

59. Bustin, S.A., Benes, V., Garson, J.A., Hellemans, J., Huggett, J., Kubista, M., Mueller, R., Nolan, T., Pfaffl, M.W., Shipley, G.L. et al. (2009) The MIQE guidelines: minimum information for publication of quantitative real-time PCR experiments. Clin Chem, 55, 611–622.

60. Pfaffl, M.W. (2001) A new mathematical model for relative quantification in real-time RT-PCR. Nucleic Acids Res, 29, e45.

61. Van Etten, J., Schagat, T.L. and Goldstrohm, A.C. (2013) A guide to design and optimization of reporter assays for 3’ untranslated region mediated regulation of mammalian messenger RNAs. Methods, 63, 110–118.

62. Bohn, J.A., Van Etten, J.L., Schagat, T.L., Bowman, B.M., McEachin, R.C., Freddolino, P.L. and Goldstrohm, A.C. (2018) Identification of diverse target RNAs that are functionally regulated by human Pumilio proteins. Nucleic Acids Res, 46, 362–386.

63. Bos, T.J., Nussbacher, J.K., Aigner, S. and Yeo, G.W. (2016) Tethered Function Assays as Tools to Elucidate the Molecular Roles of RNA-Binding Proteins. Adv Exp Med Biol, 907, 61–88.

64. Clement, S.L. and Lykke-Andersen, J. (2008) A tethering approach to study proteins that activate mRNA turnover in human cells. Methods Mol Biol, 419, 121–133.

65. Coller, J. and Wickens, M. (2007) Tethered function assays: an adaptable approach to study RNA regulatory proteins. Methods Enzymol, 429, 299–321.

66. Ozgur, S., Chekulaeva, M. and Stoecklin, G. (2010) Human Pat1b connects deadenylation with mRNA decapping and controls the assembly of processing bodies. Mol Cell Biol, 30, 4308–4323.

67. Boehm, V., Gerbracht, J.V., Marx, M.C. and Gehring, N.H. (2016) Interrogating the degradation pathways of unstable mRNAs with XRN1-resistant sequences. Nat Commun, 7, 13691.

68. Tritschler, F., Braun, J.E., Eulalio, A., Truffault, V., Izaurralde, E. and Weichenrieder, O. (2009) Structural basis for the mutually exclusive anchoring of P body components EDC3 and Tral to the DEAD box protein DDX6/Me31B. Mol Cell, 33, 661–668.

69. Brandmann, T., Fakim, H., Padamsi, Z., Youn, J.Y., Gingras, A.C., Fabian, M.R. and Jinek, M. (2018) Molecular architecture of LSM14 interactions involved in the assembly of mRNA silencing complexes. EMBO J, 37.

70. Kamenska, A., Simpson, C., Vindry, C., Broomhead, H., Benard, M., Ernoult-Lange, M., Lee, B.P., Harries, L.W., Weil, D. and Standart, N. (2016) The DDX6-4E-T interaction mediates translational repression and P-body assembly. Nucleic Acids Res, 44, 6318–6334.

71. Temme, C., Simonelig, M. and Wahle, E. (2014) Deadenylation of mRNA by the CCR4-NOT complex in Drosophila: molecular and developmental aspects. Front Genet, 5, 143.

72. Van Etten, J., Schagat, T.L., Hrit, J., Weidmann, C.A., Brumbaugh, J., Coon, J.J. and Goldstrohm, A.C. (2012) Human Pumilio proteins recruit multiple deadenylases to efficiently repress messenger RNAs. J Biol Chem, 287, 36370–36383.

73. Marzluff, W.F. and Koreski, K.P. (2017) Birth and Death of Histone mRNAs. Trends Genet, 33, 745–759.

74. Waghray, S., Williams, C., Coon, J.J. and Wickens, M. (2015) Xenopus CAF1 requires NOT1-mediated interaction with 4E-T to repress translation in vivo. RNA, 21, 1335–1345.

75. Cooke, A., Prigge, A. and Wickens, M. (2010) Translational repression by deadenylases. J Biol Chem, 285, 28506–28513.

76. Jonas, S. and Izaurralde, E. (2013) The role of disordered protein regions in the assembly of decapping complexes and RNP granules. Genes Dev, 27, 2628–2641.

77. Lykke-Andersen, J. and Wagner, E. (2005) Recruitment and activation of mRNA decay enzymes by two ARE-mediated decay activation domains in the proteins TTP and BRF-1. Genes Dev, 19, 351–361.

78. Jonas, S. and Izaurralde, E. (2015) Towards a molecular understanding of microRNA-mediated gene silencing. Nat Rev Genet, 16, 421–433.

79. Sgromo, A., Raisch, T., Bawankar, P., Bhandari, D., Chen, Y., Kuzuoglu-Ozturk, D., Weichenrieder, O. and Izaurralde, E. (2017) A CAF40-binding motif facilitates recruitment of the CCR4-NOT complex to mRNAs targeted by Drosophila Roquin. Nat Commun, 8, 14307.

80. Sgromo, A., Raisch, T., Backhaus, C., Keskeny, C., Alva, V., Weichenrieder, O. and Izaurralde, E. (2018) Drosophila Bag-of-marbles directly interacts with the CAF40 subunit of the CCR4-NOT complex to elicit repression of mRNA targets. RNA, 24, 381–395.

81. Webster, M.W., Stowell, J.A. and Passmore, L.A. (2019) RNA-binding proteins distinguish between similar sequence motifs to promote targeted deadenylation by Ccr4-Not. Elife, 8.

82. Goldstrohm, A.C., Hook, B.A., Seay, D.J. and Wickens, M. (2006) PUF proteins bind Pop2p to regulate messenger RNAs. Nat Struct Mol Biol, 13, 533–539.

83. Suh, N., Crittenden, S.L., Goldstrohm, A., Hook, B., Thompson, B., Wickens, M. and Kimble, J. (2009) FBF and its dual control of gld-1 expression in the Caenorhabditis elegans germline. Genetics, 181, 1249–1260.

84. Chen, J., Rappsilber, J., Chiang, Y.C., Russell, P., Mann, M. and Denis, C.L. (2001) Purification and characterization of the 1.0 MDa CCR4-NOT complex identifies two novel components of the complex. J Mol Biol, 314, 683–694.

85. Wagner, E., Clement, S.L. and Lykke-Andersen, J. (2007) An unconventional human Ccr4-Caf1 deadenylase complex in nuclear cajal bodies. Mol Cell Biol, 27, 1686–1695.

86. Bai, Y., Salvadore, C., Chiang, Y.C., Collart, M.A., Liu, H.Y. and Denis, C.L. (1999) The CCR4 and CAF1 proteins of the CCR4-NOT complex are physically and functionally separated from NOT2, NOT4, and NOT5. Mol Cell Biol, 19, 6642-6651.

87. Lau, N.C., Kolkman, A., van Schaik, F.M., Mulder, K.W., Pijnappel, W.W., Heck, A.J. and Timmers, H.T. (2009) Human Ccr4-Not complexes contain variable deadenylase subunits. Biochem J, 422, 443–453.

88. Goldstrohm, A.C., Seay, D.J., Hook, B.A. and Wickens, M. (2007) PUF protein-mediated deadenylation is catalyzed by Ccr4p. J Biol Chem, 282, 109–114.

89. Olivas, W. and Parker, R. (2000) The Puf3 protein is a transcript-specific regulator of mRNA degradation in yeast. Embo J, 19, 6602–6611.

90. Blewett, N.H. and Goldstrohm, A.C. (2012) A eukaryotic translation initiation factor 4E-binding protein promotes mRNA decapping and is required for PUF repression. Mol Cell Biol, 32, 4181–4194.

91. Tucker, M., Staples, R.R., Valencia-Sanchez, M.A., Muhlrad, D. and Parker, R. (2002) Ccr4p is the catalytic subunit of a Ccr4p/Pop2p/Notp mRNA deadenylase complex in Saccharomyces cerevisiae. Embo J, 21, 1427–1436.

92. Webster, M.W., Chen, Y.H., Stowell, J.A.W., Alhusaini, N., Sweet, T., Graveley, B.R., Coller, J. and Passmore, L.A. (2018) mRNA Deadenylation Is Coupled to Translation Rates by the Differential Activities of Ccr4-Not Nucleases. Mol Cell, 70, 1089–1100 e1088.

93. Yi, H., Park, J., Ha, M., Lim, J., Chang, H. and Kim, V.N. (2018) PABP Cooperates with the CCR4-NOT Complex to Promote mRNA Deadenylation and Block Precocious Decay. Mol Cell, 70, 1081–1088 e1085.

94. Thomsen, S., Anders, S., Janga, S.C., Huber, W. and Alonso, C.R. (2010) Genome-wide analysis of mRNA decay patterns during early Drosophila development. Genome Biol, 11, R93.

95. Driscoll, H.E., Muraro, N.I., He, M. and Baines, R.A. (2013) Pumilio-2 regulates translation of Nav1.6 to mediate homeostasis of membrane excitability. J Neurosci, 33, 9644–9654.

96. Sonoda, J. and Wharton, R.P. (1999) Recruitment of Nanos to hunchback mRNA by Pumilio. Genes Dev, 13, 2704–2712.

97. Naudin, C., Hattabi, A., Michelet, F., Miri-Nezhad, A., Benyoucef, A., Pflumio, F., Guillonneau, F., Fichelson, S., Vigon, I., Dusanter-Fourt, I. et al. (2017) PUMILIO/FOXP1 signaling drives expansion of hematopoietic stem/progenitor and leukemia cells. Blood, 129, 2493–2506.

98. Chen, D., Zheng, W., Lin, A., Uyhazi, K., Zhao, H. and Lin, H. (2012) Pumilio 1 Suppresses Multiple Activators of p53 to Safeguard Spermatogenesis. Curr Biol, 22, 420–425.

99. Mak, W., Fang, C., Holden, T., Dratver, M.B. and Lin, H. (2016) An Important Role of Pumilio 1 in Regulating the Development of the Mammalian Female Germline. Biol Reprod, 94, 134.

100. Zhang, M., Chen, D., Xia, J., Han, W., Cui, X., Neuenkirchen, N., Hermes, G., Sestan, N. and Lin, H. (2017) Post-transcriptional regulation of mouse neurogenesis by Pumilio proteins. Genes Dev, 31, 1354–1369.

101. Fox, M., Urano, J. and Reijo Pera, R.A. (2005) Identification and characterization of RNA sequences to which human PUMILIO-2 (PUM2) and deleted in Azoospermia-like (DAZL) bind. Genomics, 85, 92–105.

102. Moore, F.L., Jaruzelska, J., Fox, M.S., Urano, J., Firpo, M.T., Turek, P.J., Dorfman, D.M. and Pera, R.A. (2003) Human Pumilio-2 is expressed in embryonic stem cells and germ cells and interacts with DAZ (Deleted in AZoospermia) and DAZ-like proteins. Proc Natl Acad Sci U S A, 100, 538–543.

103. Siemen, H., Colas, D., Heller, H.C., Brustle, O. and Pera, R.A. (2011) Pumilio-2 function in the mouse nervous system. PLoS One, 6, e25932.

104. Urano, J., Fox, M.S. and Reijo Pera, R.A. (2005) Interaction of the conserved meiotic regulators, BOULE (BOL) and PUMILIO-2 (PUM2). Mol Reprod Dev, 71, 290-298.

105. Xu, E.Y., Chang, R., Salmon, N.A. and Reijo Pera, R.A. (2007) A gene trap mutation of a murine homolog of the Drosophila stem cell factor Pumilio results in smaller testes but does not affect litter size or fertility. Mol Reprod Dev, 74, 912–921.

106. Vessey, J.P., Schoderboeck, L., Gingl, E., Luzi, E., Riefler, J., Di Leva, F., Karra, D., Thomas, S., Kiebler, M.A. and Macchi, P. (2010) Mammalian Pumilio 2 regulates dendrite morphogenesis and synaptic function. Proc Natl Acad Sci U S A, 107, 3222–3227.

107. Gennarino, V.A., Singh, R.K., White, J.J., De Maio, A., Han, K., Kim, J.Y., Jafar-Nejad, P., di Ronza, A., Kang, H., Sayegh, L.S. et al. (2015) Pumilio1 haploinsufficiency leads to SCA1-like neurodegeneration by increasing wild-type Ataxin1 levels. Cell, 160, 1087–1098.

108. Gennarino, V.A., Palmer, E.E., McDonell, L.M., Wang, L., Adamski, C.J., Koire, A., See, L., Chen, C.A., Schaaf, C.P., Rosenfeld, J.A. et al. (2018) A Mild PUM1 Mutation Is Associated with Adult-Onset Ataxia, whereas Haploinsufficiency Causes Developmental Delay and Seizures. Cell, 172, 924–936 e911.

109. Miles, W.O., Tschop, K., Herr, A., Ji, J.Y. and Dyson, N.J. (2012) Pumilio facilitates miRNA regulation of the E2F3 oncogene. Genes Dev, 26, 356–368.

110. Follwaczny, P., Schieweck, R., Riedemann, T., Demleitner, A., Straub, T., Klemm, A.H., Bilban, M., Sutor, B., Popper, B. and Kiebler, M.A. (2017) Pumilio2-deficient mice show a predisposition for epilepsy. Dis Model Mech, 10, 1333–1342.

111. Wu, X.L., Huang, H., Huang, Y.Y., Yuan, J.X., Zhou, X. and Chen, Y.M. (2015) Reduced Pumilio-2 expression in patients with temporal lobe epilepsy and in the lithium-pilocarpine induced epilepsy rat model. Epilepsy Behav, 50, 31–39.

112. Long, E.O. and Dawid, I.B. (1980) Alternative pathways in the processing of ribosomal RNA precursor in Drosophila melanogaster. J Mol Biol, 138, 873–878.

113. Tautz, D., Hancock, J.M., Webb, D.A., Tautz, C. and Dover, G.A. (1988) Complete sequences of the rRNA genes of Drosophila melanogaster. Mol Biol Evol, 5, 366–376.

